# A Modular Bacteriophage T4 Nanoparticle Platform Enables Rapid Design of Dual COVID-19-Flu Mucosal Vaccines

**DOI:** 10.1101/2024.10.09.617418

**Authors:** Jingen Zhu, Jian Sha, Himanshu Batra, Swati Jain, Xiaorong Wu, Emily K. Hendrix, Paul B. Kilgore, Keer Sun, Kenneth S. Plante, Jessica A. Plante, Jordyn Walker, Pan Tao, Ashok K. Chopra, Venigalla B. Rao

**Affiliations:** Bacteriophage Medical Research Center, Department of Biology, The Catholic University of America, Washington, DC, 20064, USA; Department of Microbiology and Immunology, University of Texas Medical Branch, Galveston, TX, 77555, USA; Galveston National Laboratory, University of Texas Medical Branch, Galveston, TX, 77555, USA; Institute for Human Infections and Immunity, University of Texas Medical Branch, Galveston, TX, 77555, USA; World Reference Center for Emerging Viruses and Arboviruses, University of Texas Medical Branch, Galveston, TX, 77555, USA; Center for Biodefense and Emerging Infectious Diseases, University of Texas Medical Branch, Galveston, TX, 77555, USA; Sealy Institute for Vaccine Sciences, University of Texas Medical Branch, Galveston, TX, 77555, USA; State Key Laboratory of Agricultural Microbiology, College of Veterinary Medicine, Huazhong Agricultural University, Wuhan, Hubei, China

## Abstract

A multivalent, rapidly deployable, mucosal vaccine platform is desperately needed to prevent the acquisition and transmission of respiratory infections during epidemics and pandemics. We present one such bacteriophage T4-based platform, and design of dual COVID-19-Flu mucosal vaccines by exploiting its unique architecture. These include: T4’s natural affinity for nasal mucosa, flexible engineering to incorporate multiple antigens, and repeat and symmetric epitope presentation for enhanced B cell responses. Hundreds of SARS-CoV-2 spike trimers and nucleocapsid proteins, and influenza hemagglutinin trimers and M2e peptides, were incorporated into a single phage, creating the highest density nanoparticle presentation yet reported. Intranasal administration of adjuvant-free vaccine induced robust mucosal immunity in mice including, neutralizing antibody and secretory IgA, lung-resident CD4^+^/CD8^+^ T cells, diverse memory B cells, and complete protection against SARS-CoV-2 and influenza challenges. The noninfectious T4 phage offers an extraordinary platform to rapidly design potent mucosal vaccines against emerging bacterial and viral threats.

## Introduction

It is now abundantly clear that new and innovative vaccine design strategies are desperately needed to address future epidemics and pandemics. As effective as the current vaccine platforms are, ∼65% of people in low- and middle-income communities are yet to receive a single dose of the COVID-19 vaccine. The current vaccines are not thermostable and/or expensive to manufacture and distribute to many global communities. Of particular urgency is the need for a rapidly deployable mucosal vaccine platform that can curb respiratory infections at the portal of entry and minimize person-to-person transmission^1^.

A case in point is the continuing COVID-19 pandemic, caused by severe acute respiratory syndrome coronavirus 2 (SARS-CoV-2), which had a devastating impact on global health claiming over 7 million lives since 2019 (World Health Organization [WHO], https://covid19.who.int). Simultaneously, seasonal influenza (Flu) epidemics, caused by influenza A and B viruses, remain a major threat to global health, causing up to ∼650,000 deaths annually^2^. Recent outbreaks of highly pathogenic avian influenza and the current circulating strains in farm animals and humans, further raise concerns about the potential emergence of a novel pandemic strain (Centers for Disease Control and Prevention [CDC], https://cdc.gov/bird-flu/). Co-circulation of SARS-CoV-2 and influenza viruses is another cause for alarm, as both pathogens share similar transmission routes, target tissues, and seasonal patterns, and can lead to more severe disease outcomes in co-infected individuals^3–6^. While injectable vaccines such as the ones based on mRNA and adenoviral vectors are highly effective in minimizing severe disease^7,8^, they do not induce sufficient mucosal immunity in the respiratory tract to prevent infection or transmission^1,9–11^. Compounding this problem is the antigenic drift and shift by influenza or SARS-CoV-2 variants requiring repeated and yearly revaccinations to generate protection against circulating strains^12^-^14^.sub>

There are three potential strategies to address these challenges. First is to design intranasal mucosal vaccines that can induce robust mucosal immunity in addition to systemic responses and provide a first line of defense at the point of entry^15,16^. Second is to have a flexible engineering platform that allows incorporation of multiple antigen targets into a vaccine formulation to generate broad protection against diverse variants^17–19^. Third is high density and systematic epitope presentation on a nanoparticle surface for high avidity and B cell receptor cross-linking as well as recruitment of low-affinity B cells for greater B cell clonotype diversity and breadth of neutralization_20-23._

Bacteriophages have emerged as promising VLP (virus-like particle) vaccine design platforms due to their ordered antigen display that enhances immune recognition, cost-effective production in bacteria, and exceptional stability, making them particularly attractive for global vaccine development^24–29^. Here, we present a pinnacle of VLP vaccine designs that integrates all the above-desired strategies into one platform by exploiting the unique architecture of bacteriophage (phage) T4^30–32^. Indeed, we pushed the boundaries of the T4 nanoparticle platform by incorporating five distinct and conserved antigen targets from two deadly respiratory viruses, SARS-CoV-2 and influenza, to create a highly efficacious dual-pathogen mucosal vaccine. The antigens include: spike ectodomain (S-ecto) and nucleocapsid protein (NP) of SARS-CoV-2^33^, hemagglutinin stem domain (HA-stem)^34,35^, the extracellular domain of the matrix protein 2 (M2e), and matrix protein 1 (M1) of influenza^13,36^. These relatively well-conserved antigens but highly diverse in structure and function that have never been combined in a vaccine platform are selected here to test the limits of the platform, and its ability to induce multi-pronged immunity; humoral, cellular, and mucosal.

We first hard-wired the T4 phage by inserting M2e-Hoc, Soc-SpyCatcher, and NP or M1 genes under the control of phage promoters into the phage genome by *in vivo* CRISPR-engineering^37–39^. The recombinant phages thus produced display arrays of M2e and SpyCatcher molecules as fusions of the T4 outer capsid proteins Hoc (highly antigenic outer capsid protein) and Soc (small outer capsid protein), respectively, on its large 120 x 86 nm capsid^40–42^, while the NP and M1 molecules were packaged in the capsid interior by virtue of their fusion to a capsid targeting sequence (CTS)^43,44^. SpyTagged S-ecto and HA-stem trimers were then covalently conjugated *in vitro* to the surface-displayed SpyCatcher molecules^45^. Hundreds of target antigen molecules incorporated into different structural compartments generated a highly decorated phage making it the highest-density nanoparticle presentation reported to date. Yet, the process is quite simple and can be rapidly applied to any respiratory pathogen to produce a series of vaccine candidates in a matter of weeks as was accomplished in this study.

Phage T4 exhibits natural affinity to mucin glycoproteins secreted by mucosal epithelia^46,47^. The binding of 180Å-long Hoc fibers containing three Ig-like domains to mucins leads to the persistence of T4 vaccines in the upper and lower respiratory tracks including the lungs for at least 4-weeks^48^, allowing efficient antigen presentation and continuous stimulation of the mucosal immune system^48,49^. Consequently, intranasal administration of two doses of T4-COVID-Flu vaccine constructed as above induced robust mucosal immunity against both SARS-CoV-2 and influenza infections, in addition to eliciting strong humoral and cellular responses. Remarkably, the mucosal responses included high titers of secretory IgA in serum and lung fluids as well as induction of effector and memory T cell responses and lung-resident CD4^+^ and CD8^+^ T cells. Furthermore, the vaccine-induced strong virus neutralization titers, balanced Th1 and Th2 responses, diverse memory B cell populations, as well as complete protection against lethal challenges with the prototypic SARS-CoV-2 and influenza viruses.

The body of evidence demonstrates that T4 possesses characteristics of an ideal global mucosal vaccine design platform encapsulating all the key desirable features in one vehicle. These include: non-replicating bio-vector, plug-and-play multivalent design, adaptability to diverse antigens, simple vaccine formulation in phosphate buffer-saline (PBS) free of adjuvants and chemicals, thermostability, and inexpensive scalability and manufacturability, thus making it a powerful protein-based mucosal vaccine design platform to meet the demands of a countermeasure against future epidemic and pandemic emerging bacterial and viral threats.

## Results

### Overall design of T4-CoV-Flu dual vaccine candidates

Our overall design was focused on creating T4-CoV-Flu dual vaccine candidates using the conserved and structural elements of both SARS-CoV-2 and influenza viruses. Taking advantage of the unique architectural features of the phage T4 platform, we set out to strategically incorporate several antigen targets into the T4 nanoparticle structure (Fig. 1; Supplementary Fig 1a, b). These include: (1) the mammalian cell-expressed and prefusion-stabilized S-ecto trimers and HA-stem trimers, which would be displayed on the capsid exterior using the SpyCatcher-Spytag conjugation *in vitro*^45^, (2) the 24-amino acid (aa) influenza M2e peptide fused to the tips of the 180 Å-long Hoc fibers, which would be displayed through *in vivo* assembly, and (3) the conserved SARS-CoV-2 NP or influenza M1 proteins, which would be packaged *in vivo* in the interior of the capsid.

**Fig. 1:**
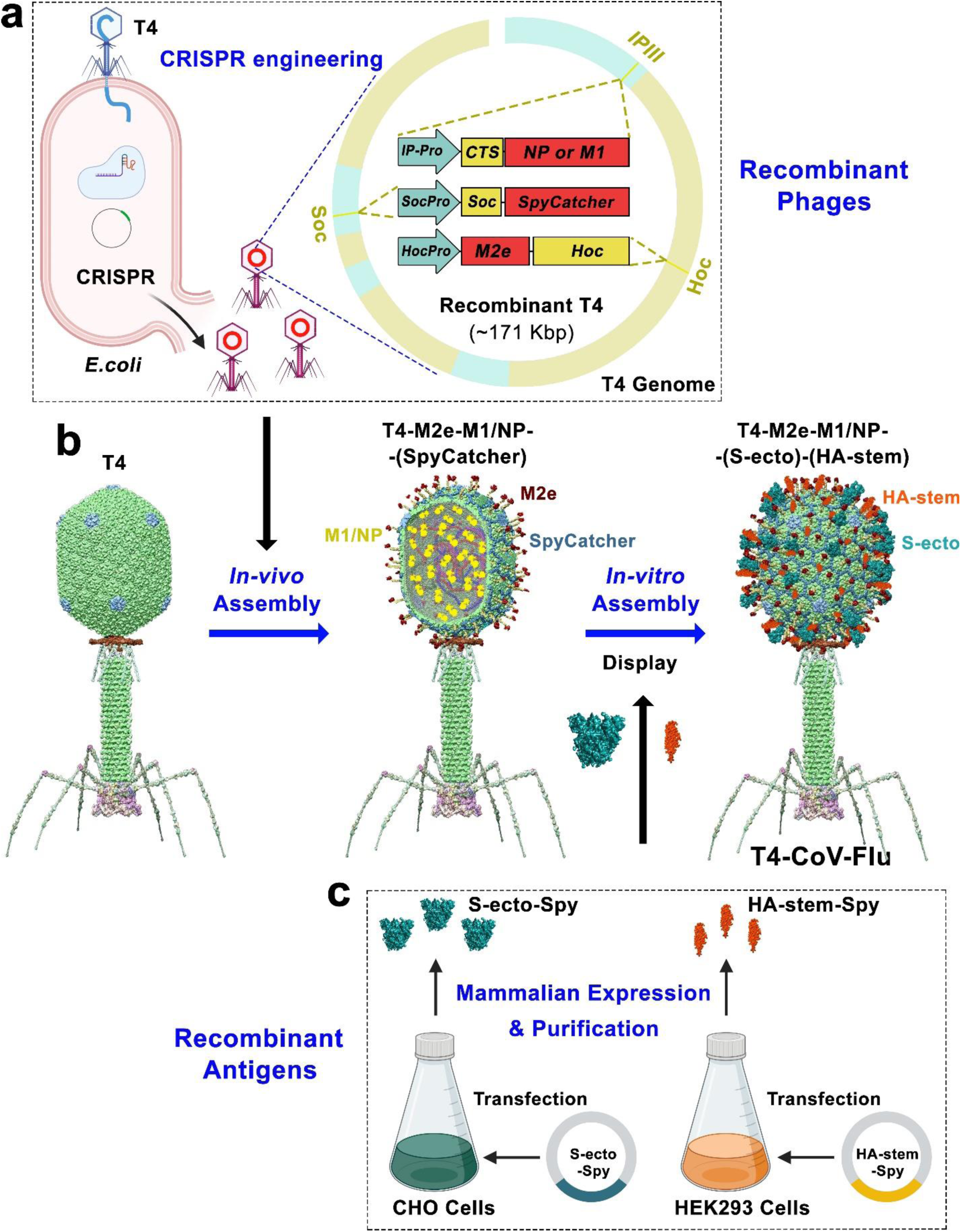
A two-step strategy to design phage T4-CoV-Flu multivalent vaccine. **a** CRISPR engineering of recombinant T4 phages displaying Flu-M1, Flu-M2e, CoV-NP, and SpyCatcher in *E. coli*. **b** Assembly of T4-CoV-Flu vaccine through *in vivo* antigen display and *in vitro* SpyCatcher-SpyTag conjugation of mammalian cell-expressed S-ecto and HA-stem trimers. **c** Production of S-ecto-Spy and HA-stem-Spy trimers in CHO and HEK293F cells for phage conjugation.

*In vitro* assembly was used for displaying the envelope trimers because doing so *in vivo* during phage infection in *E. coli* was proven to be challenging for several reasons, such as poor expression and folding, and lack of post-translational modifications. Therefore, the mammalian cell-expressed proteins were biochemically characterized and displayed on the capsid through *in vitro* assembly, which also ensured the structural and functional integrity of the antigen epitopes (Fig. 1b, c; Supplementary Fig. 1c).

On the other hand, the M2e, NP, and M1 antigens requiring no essential post-translational modifications were well-expressed during phage infection when the target genes were inserted into the T4 genome by CRISPR genome engineering under the control of strong phage promoters. The expressed proteins were then either displayed on the capsid due to their fusion to Hoc or Soc, or packaged inside the capsid due to their fusion to T4 scaffolding core-targeting CTS sequence (Fig. 1a, b).

To maximize expression, we employed codon optimization because significant differences in nucleotide content exist between the T4 phage genome (65% AT + 35% GC) and the *E. coli* host genome (49% AT + 51% GC) (Fig. 2a). Comparison of display efficiencies showed that the T4 codon-optimized Soc-SpyCatcher exhibited significantly higher display amounts (Fig. 2b, c; 205 copies per capsid) and correspondingly higher conjugation efficiency of SpyTagged trimers (Fig. 2d, e; see below). Notably, such hard-wired SpyCatcher recombinant phage provided a large surface lattice to which any pathogen antigen or a mixture of pathogens could be tethered to generate multivalent vaccines.

**Fig. 2:**
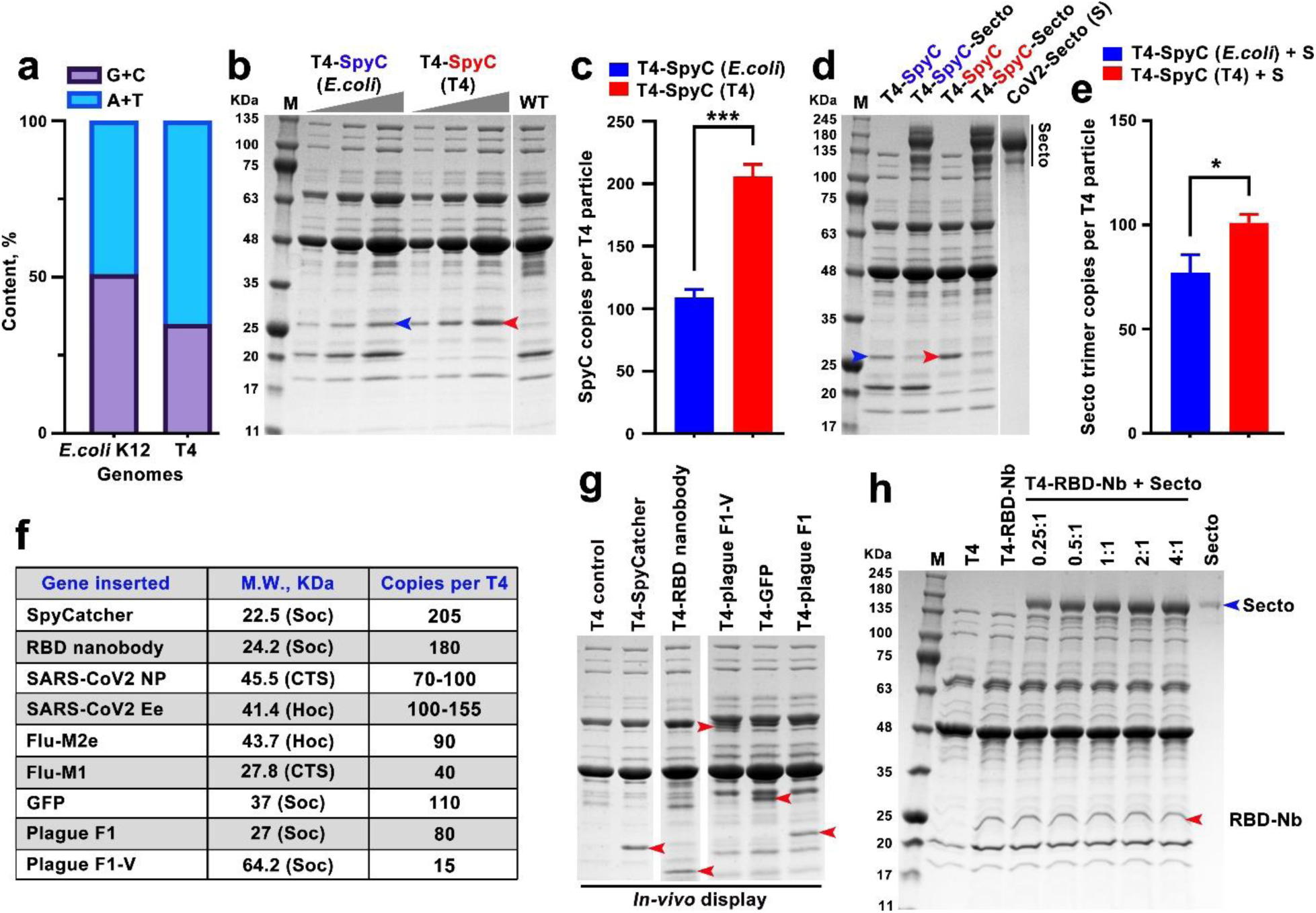
Optimized *in vivo* display and encapsulation of multiple pathogen antigens into T4 nanoparticle. **a** Comparison of nucleotide content (A+T and G+C) in *E. coli* K12 and T4 genomes. **b** and **c** Comparison of display efficiency between *E. coli*-codon (blue) and T4-codon (red) optimized Soc-SpyCatcher (SpyC) by SDS–polyacrylamide gel electrophoresis (PAGE) (b) and quantitative analysis (c). The molecular weight standards (M) in KDa are shown on the left of the gel. **d** and **e** SDS-PAGE (d) and quantitative analysis (e) of Spy-tagged S-ecto trimers displayed on the T4 capsid. The interaction efficiency between S-ecto trimers and *E. coli*- or T4-codon-optimized SpyC was assessed, demonstrating improved display with T4-codon-optimized SpyC. **f** Table showing various antigens of interest either displayed on or encapsidated into the T4 capsid, their sizes, and copy numbers. **g** SDS-PAGE showing the display of representative proteins on the T4 capsid. Red arrowheads indicate the positions of the corresponding proteins. **h** SDS-PAGE showing display of S-ecto trimers on T4 capsid through S-ecto and RBD nanobody (RBD-Nb) interaction. The positions of S-ecto and RBD-Nb are indicated. A nonparametric Student’s *t*-test was used to compare T4-SpyC (*E. coli*) versus T4-SpyC (T4) groups in (c) and (e). *, P < 0.05; ***, P < 0.001. Data represent means ± standard deviation (SD) from three replicates.

Taking advantage of these features, we performed extensive series of experiments to optimize the T4 platform for efficient *in vivo* display on the capsid surface and/or encapsulation in the capsid interior using diverse antigens from SARS-CoV-2 and influenza, as well as several other model antigens, nanobodies, and fluorescent proteins (Fig. 2, f to h). Notably, we achieved display of up to 180 copies of functional nanobodies per capsid, demonstrating the platform’s potential for applications beyond traditional vaccines. These optimizations were crucial for enabling the efficient production of our multivalent T4-CoV-Flu dual vaccine and highlight the broader potential of the T4 platform for rapid vaccine development against various pathogens.

Finally, a series of vaccine candidates containing combinations of SARS-CoV-2 and Flu antigens were rapidly produced and tested in a mouse model that allowed the selection of a candidate that showed the highest efficacy as a dual CoV-Flu vaccine.

### Production of T4-CoV-Flu vaccine candidates

The T4-CoV-Flu vaccine candidates were produced by a two-step process; first engineer recombinant phages containing displayed or encapsidated antigens by *in vivo* expression, then perform *in vitro* assembly and conjugation using purified mammalian cell-expressed antigens (Fig. 1). For the former, gene segments encoding influenza M1 (or NP), M2e, and SpyCatcher were inserted into specific loci of the T4 genome; CTS-IP, Hoc, and Soc, respectively, using CRISPR-targeted genome editing (Supplementary Fig. 1a, b). SDS-PAGE and Western blot analyses of CsCl-gradient purified recombinant phage particles showed successful incorporation and assembly of each of these antigen components (Fig. 3a). Molecular weight determinations further confirmed M2e-Hoc fusion, Soc-SpyCatcher conjugation to SpyTagged S-ecto and HA-stem trimers, and packaging of M1 and NP proteins.

**Fig. 3:**
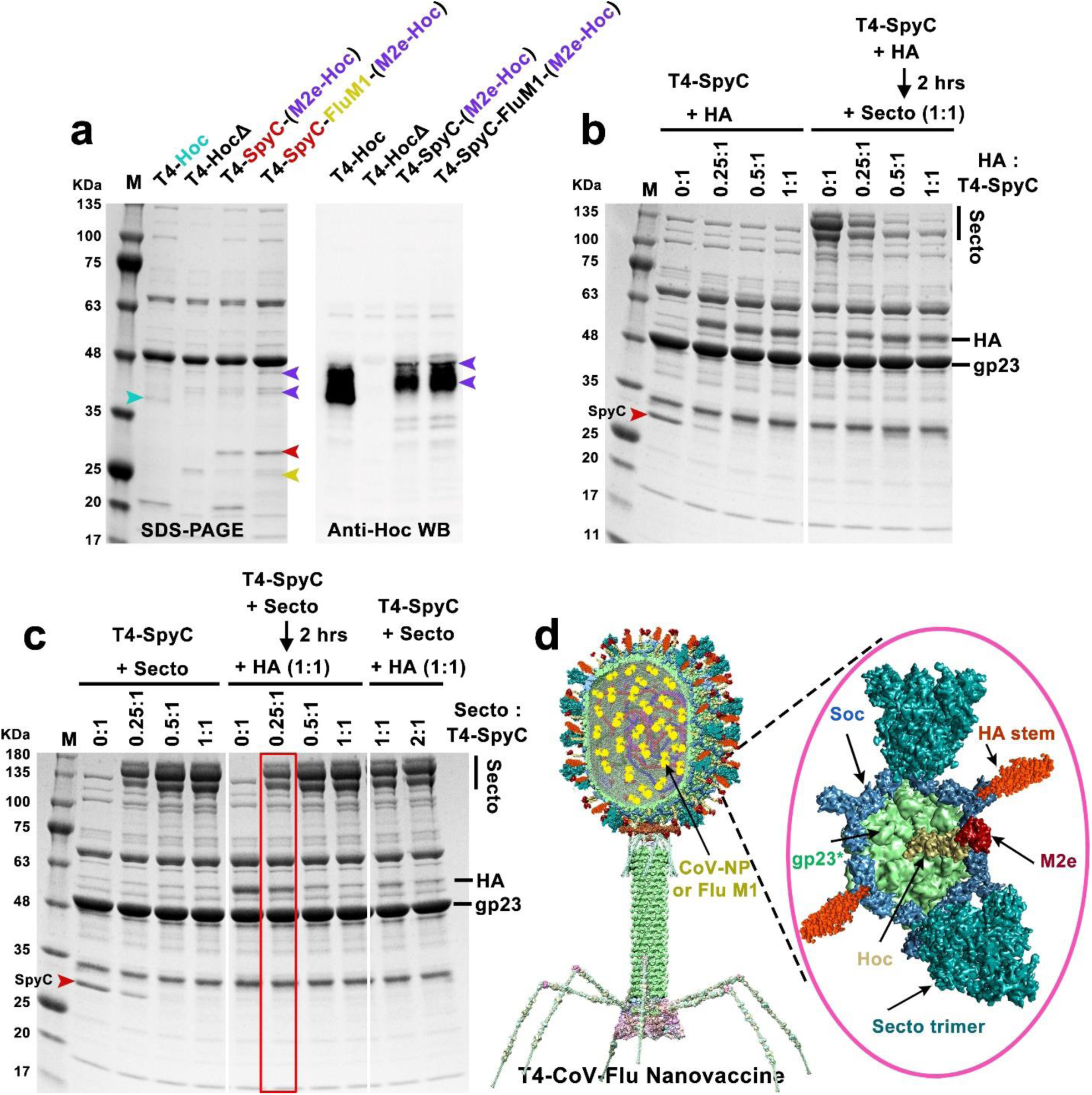
Assembly of the multivalent T4-CoV-Flu nanovaccine. **a** SDS-PAGE and Western Blot (WB) analyses of phage nanoparticles displaying M2e and SpyC and encapsulating M1 *in vivo*. Note that due to the similar molecular size of Hoc and M2e-Hoc to other T4 structural proteins, a Hoc-specific antibody was used to detect Hoc and M2e-Hoc. Arrowheads with various colors indicate the corresponding displayed or encapsidated antigens. **b** and **c** Optimization of the co-display of S-ecto and HA-stem trimers. (b) *In vitro* display of HA stem trimers on T4-SpyC phage (left) and sequential co-display of HA stem and S-ecto trimers (HA added first, incubated, then S-ecto) (right) at increasing ratios of HA stem trimer molecules to Soc binding sites (0:1 to 1:1). The ratio of S-ecto trimer molecules to Soc binding sites was fixed at 1:1 in the co-display assay. (c) *In vitro* display of S-ecto trimers on T4-SpyC phage (left), sequential co-display of S-ecto and HA stem trimers (S-ecto added first, incubated, then HA) (middle), and simultaneous co-display of S-ecto and HA stem trimers (both added at the same time) (right) at increasing ratios of S-ecto trimer molecules to Soc binding sites (0:1 to 2:1). The ratio of HA stem trimer molecules to Soc binding sites was fixed at 1:1 in the co-display assay. **d** Structural model of the final T4-CoV-Flu nanovaccine. Inset: Enlarged view of a single hexameric capsomer showing six subunits of major capsid protein gp23* (green) associated with trimers of Soc (blue) and Hoc fiber (yellow). The Flu M2e peptide (dark red) is displayed at the tips of Hoc fibers, while the SpyCatcher-Soc fusions surround the capsomer periphery. The CoV NP or Flu M1 molecules (yellow) are packaged inside the capsid. Spy-tagged S-ecto trimers (cyan) and HA stem trimers (red) are co-conjugated to Soc-SpyCatcher trimers.

Approximately 40 copies of M1 or 100 copies of NP, 90 copies of M2e, and 205 copies of Soc-SpyCatcher were co-incorporated into a single T4 nanoparticle (Fig. 3a). It is noteworthy that such a multi-antigen *in vivo* engineering of phage has not been reported so far, pointing to the versatility of the T4 vaccine design platform. Notably, incorporation of these antigens did not affect phage yield, which was in the order of 10^14^-10^15^ phage particles per liter of *E. coli* culture. Assuming about 10^11^-10^12^ particles per vaccine dose, this reflects up to ∼10^3^ to 10^4^ doses per liter of *E. coli*, which might be further improved in an industrial-scale fermenter under optimal nutrient and aeration conditions.

In the second step, S-ecto and HA-stem envelope trimers containing C-terminal SpyTag fusions were expressed and purified from Chinese hamster ovary (CHO) cells and human embryonic kidney 293 F (HEK 293F) cells, respectively (Supplementary Fig. 1c), and conjugated *in vitro* to Soc-SpyCatcher arrayed on T4 capsid. Both S-ecto and HA stem trimers can be efficiently conjugated to the capsid, reaching saturation at a molar ratio of 1:1 of trimer to Soc binding sites, resulting in approximately 118 copies of S-ecto trimer or 107 copies of HA-stem trimer per capsid (Supplementary Fig. 2). Since Soc-SpyCatcher subunits are located at the quasi-3-fold axes of the T4 icosahedral structure, the conjugated trimers would occupy symmetric positions on the nanoparticle surface.

To achieve optimal antigen density and ensure efficient co-display of both S-ecto and HA-stem trimers on the same nanoparticle, the ratios of SpyTagged HA-stem and S-ecto trimers to SpyCatcher sites were systematically titrated (Fig. 3b, c). The data showed that the display efficiency and copy number could be tuned by adjusting the ratios of trimers to binding sites and/or the sequence of addition of the two types of trimers. Three different sequences were explored (Fig. 3b, c): 1) HA-stem added first, incubated, and then S-ecto added; 2) S-ecto added first, incubated, and then HA-stem added; and 3) both HA-stem and S-ecto added simultaneously while varying the ratios.

The results showed approximately equivalent co-display of S-ecto and HA-stem trimers on a single capsid (∼56 copies of S-ecto trimer and ∼46 copies of HA-stem trimer) when S-ecto trimers were added first at a 0.25:1 ratio, incubated, and then HA-stem trimers were added at a 1:1 ratio. Not surprisingly, the major size difference between the S-ecto trimer (433.5 kDa) and the HA-stem trimer (100 kDa) affected the display efficiency. Displaying first the 4.3-times larger S-ecto trimers left a significant fraction of binding sites unoccupied probably due to steric hindrance, which were then filled by the much smaller HA-stem trimers (46 copies). The maximum copy number attained was ∼100 trimers per capsid. In the opposite sequence, when HA-stem trimers were added first, fewer binding sites were left unoccupied for the much larger S-ecto trimers leading to ∼35 copies per capsid (Fig. 3b, c).

The above two-step process leverages the versatility of both the CRISPR-based *in vivo* engineering and SpyCatcher-Spytag-based *in vitro* conjugation, enabling the incorporation of multiple desired antigens into a single nanoparticle at high copy numbers (Fig. 3d). Furthermore, the ability to generate desired combination of antigens and tuning their copy numbers highlight the flexibility and modularity of the phage T4 vaccine platform.

### Intranasal administration of T4-CoV-Flu vaccine candidates elicited robust systemic humoral immune responses

To evaluate the immunogenicity of the T4-CoV-Flu vaccine candidates, the recombinant phages were purified by two CsCl-gradient centrifugations and suspended in simple phosphate-buffer-saline (PBS, pH 7.4) with no adjuvant or stabilizing chemicals. Such preparations contained low levels of endotoxin (∼10 endotoxin units per 10^11^ phage), well below the FDA-recommended levels^50^. The ability of these phages to induce humoral antibody responses against various SARS-CoV-2 and influenza antigens was tested in the BALB/c mouse model by intranasal administration (Fig. 4a).

**Fig. 4:**
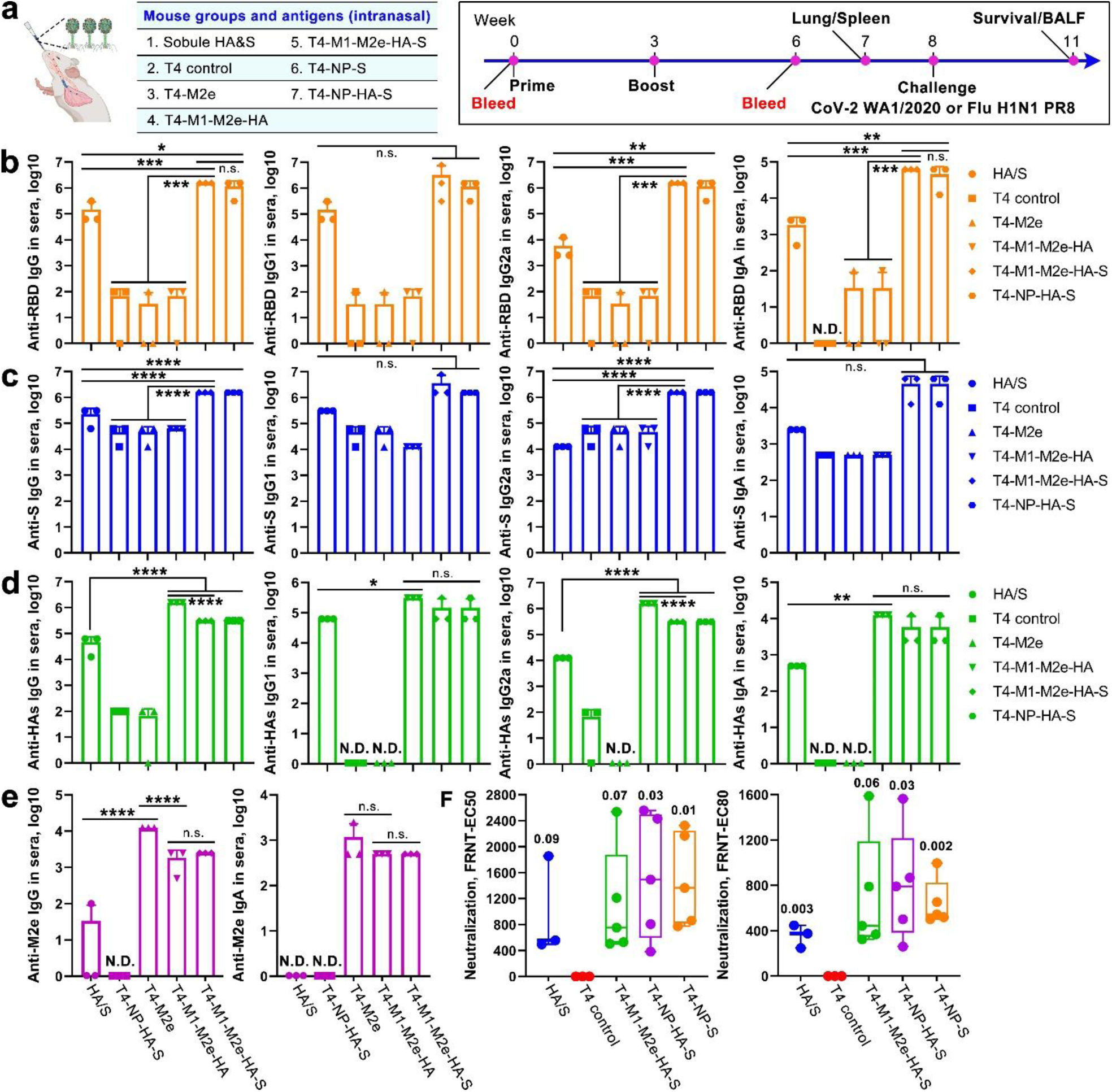
Intranasal immunization with T4-CoV-Flu vaccine elicited strong systemic humoral immune responses against multiple SARS-CoV-2 and influenza antigens. **a** Schematic representation of intranasal mouse vaccinations with various T4-CoV-Flu vaccine candidates. **b** to **e** Antibody responses against SARS-CoV-2 and influenza antigens in the sera of immunized BALB/c mice at day 21 post-final dose. ELISA was used to measure the reciprocal endpoint antibody titers for anti-RBD IgG/IgG1/IgG2a/IgA (b), anti-S-ecto IgG/IgG1/IgG2a/IgA (c), anti-HA IgG/IgG1/IgG2a/IgA (d), and anti-M2e IgG/IgA (e). **f** Serum neutralizing activity measured by Focus Reduction Neutralization Test (FRNT) assay using live SARS-CoV-2 US-WA-1/2020 strain. FRNT EC_50_ and EC_80_ values for individual serum samples from control and vaccine groups are shown. Data represent means ± SD from three pooled independent experiments (n=5). N.D. indicates not detected. Statistical comparisons among multiple groups in (b) to (e) were made using one-way analysis of variance (ANOVA) with Tukey’s post *hoc* test. *, *P* < 0.05; **, *P* < 0.01; ***, *P* < 0.001; ****, *P* < 0.0001; n.s., not significant. In (f), Student’s *t*-test was applied to compare HA/S and T4 CoV-Flu vaccine groups with T4 control. *P* values are indicated.

Groups of mice received two intranasal doses of 2.5 x 10^11^ phage particles of various vaccine candidates; T4-CoV-NP-S-ecto, T4-Flu-M2e, T4-Flu-M1-M2e-HA-stem, T4-CoV-NP-S-ecto/Flu-HA-stem, or T4-CoV-S-ecto/Flu-M1-M2e-HA-stem. Mice receiving T4 phage backbone lacking any CoV or Flu antigens were used as negative controls, while a mixture of soluble S-ecto and HA-stem trimers served as a positive control. In a previous studies, intranasal vaccination of unadjuvanted CoV-2 S-ecto or Flu HA protein was reported to induce protective mucosal immunity^51,52^.

All the T4-CoV-Flu candidates elicited high titers of serum antibodies against the externally displayed antigen components, including S-ecto, HA-stem, and M2e (Fig. 4, b to e; Supplementary Fig. 3a, b). The T4 vector-specific antibodies for reasons unknown mildly cross-reacted with S-ecto trimer. Hence, RBD which exhibits no significant cross-reactivity with T4 antibodies was also included in our analyses. The endpoint IgG titers against RBD and HA-stem trimers were at the order of 10^5^-10^6^, ∼10-fold higher than that induced by soluble trimers. Unlike the soluble trimers which bias the response to IgG1 antibodies, the T4-CoV-Flu vaccine candidates generated high and balanced levels of both IgG1 and IgG2a responses.

Interestingly, the HA-specific antibody titers in the high occupancy HA-stem display group (T4-M1-M2e-HA) were approximately 5-fold higher than the titers in the half-amount HA-stem display group (T4-M1-M2e-HA-S or T4-NP-HA-S) (Fig. 4d), indicating a dose-dependent response. Notably, compared to soluble trimers and recombinant subunit vaccines in general that show a strong bias towards inducing type 2 helper (Th2) antibody responses, the T4-CoV-Flu vaccine candidates induced high levels of both IgG2a and IgG1 responses (Supplementary Fig. 3c, d). Such a balanced Th1/Th2 antibody profile is desirable for minimizing the risk of vaccine-enhanced respiratory disease^53,54^. The Th1/Th2 balanced response may be attributed to the intrinsic adjuvant properties of the T4 phage nanoparticle, which can stimulate innate immune responses and promote antigen presentation by antigen-presenting cells^30,32,49,55^.

In addition to the antibodies targeting the displayed antigens, IgG responses were also detected against the internally packaged NP and M1 proteins, but at much lower levels (Supplementary Fig. 4). This means that the encapsidated antigens unlike the well-exposed surface epitopes were not well presented to the immune system for eliciting humoral responses. Remarkably, the surface exposed S-ecto and HA-stem antigens induced potent serum IgA responses with endpoint titers ranging from 10^4^ - 10^5^, ∼50-100 fold higher than that induced by the soluble trimers (Fig. 4, b to e). High titers of IgA antibodies are considered desirable for the clinical efficacy of vaccines because they are reported to possess anti-inflammatory activity, and are more effective than IgG in neutralizing the SARS-CoV-2 virus in the early stages of infection^56^. Moreover, IgA is the predominant antibody isotype at mucosal surfaces and plays a crucial role in protecting against transmission of respiratory pathogens^57^.

To independently evaluate the observed high immunogenicity of T4-CoV-Flu vaccine candidates, we immunized transgenic mice expressing the human angiotensin-converting enzyme 2 receptor (hACE2), the primary entry receptor for SARS-CoV-2. Consistent with the results described above in BALB/c mice, the T4-CoV-Flu vaccine candidates elicited similarly potent antibody responses against S-ecto and HA antigens in hACE2-transgenic AC70 mice (Supplementary Fig. 5), indicating the effectiveness of this vaccine strategy in an animal model that more closely mimics human infection. Importantly, these antibodies also showed strong virus-neutralizing activity as determined by the Vero E6 cell Focus Reduction Neutralization Test (FRNT) assay using the live SARS-CoV-2 US-WA-1/2020 strain (Fig. 4f), underscoring the potential of these T4-based vaccines to confer protective immunity.

Collectively, the above datasets showed robust and simultaneous induction of antibodies against two distinct respiratory viral pathogens, highlighting the potential of intranasal T4 immunization for multivalent vaccine development.

### T4-CoV-Flu vaccines induced strong mucosal humoral immune responses

To assess if the T4-CoV-Flu vaccines elicited mucosal antibody responses, we collected bronchoalveolar lavage fluid (BALF) and determined the antigen-specific antibody titers by ELISA (Fig. 4). Remarkably, all the T4-CoV-Flu phages elicited high titers of IgG, both IgG1 and IgG2a subclasses, and secretory IgA (sIgA) in BALF against the displayed antigens, including RBD/S-ecto, HA-stem, and M2e (Fig. 5). The mucosal IgG included balanced Th1-biased IgG2a and Th2-biased IgG1 subtype antibodies (Fig. 5), consistent with the balanced Th1/Th2 profile observed in the systemic antibody responses (Fig. 4). Moreover, the mucosal antibody titers induced by the T4-CoV-Flu phage nanoparticles were 5- to 25-fold higher than those induced by soluble trimers. This again highlights the effectiveness of T4-vectored mucosal delivery in stimulating local antibody responses without the need for additional adjuvants. The induction of high-titer mucosal IgG and sIgA antibodies is crucial for neutralizing the invading respiratory viruses at the portal of entry, reducing the risk of infection, disease severity, and transmission^56,58^.

**Fig. 5:**
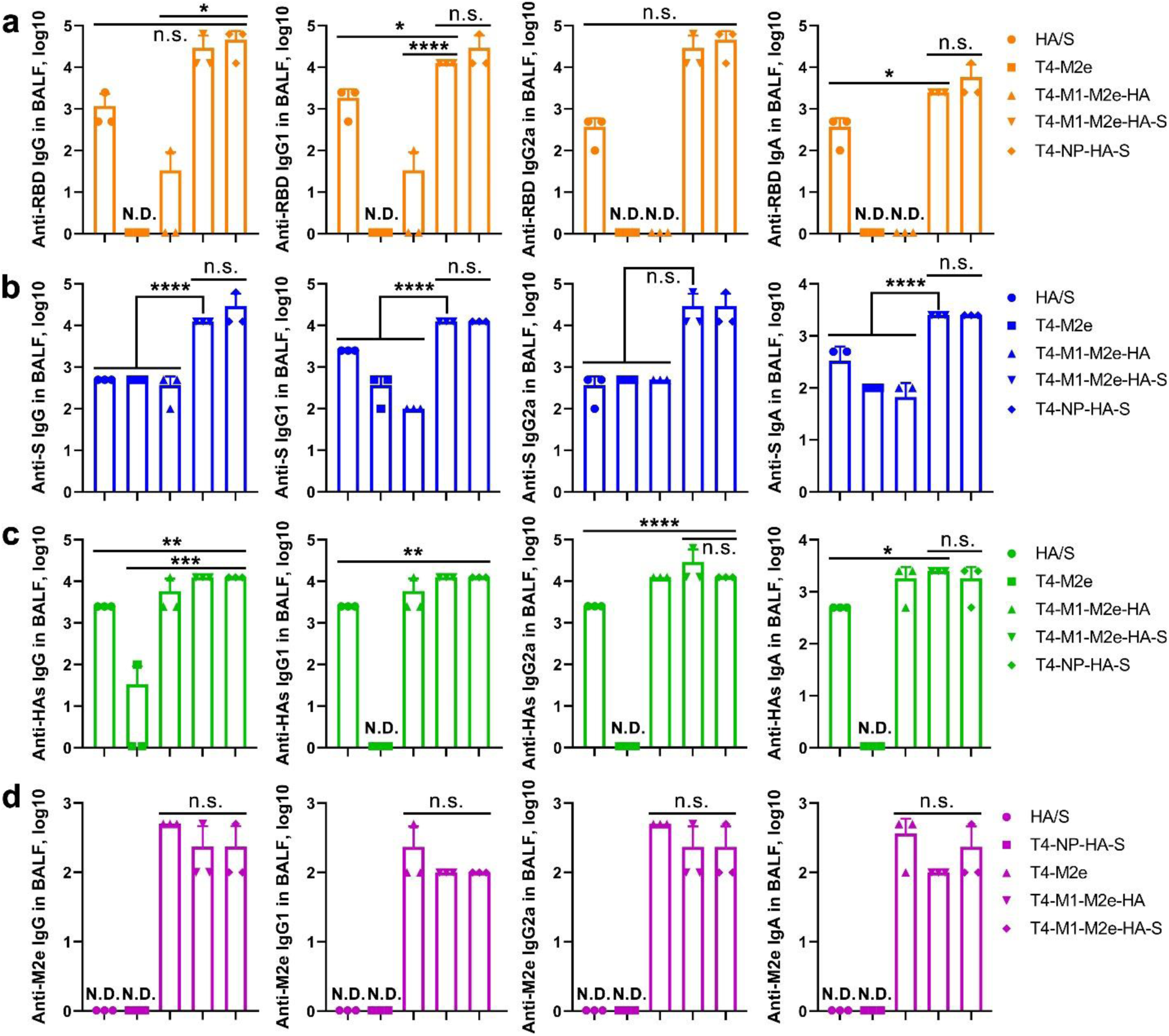
The T4-CoV-Flu intranasal vaccine induced robust mucosal humoral immune responses. The reciprocal endpoint antibody titers in BALF of anti-RBD IgG/IgG1/IgG2a/IgA (**a**), anti-S-ecto IgG/IgG1/IgG2a/IgA (**b**), anti-HA IgG/IgG1/IgG2a/IgA (**c**), and anti-M2e IgG/IgG1/IgG2a/IgA (**d**) are shown. Data represent means ± SD. N.D., not detected. The data are from three pooled independent experiments (n=5). Statistical comparisons among multiple groups were made using one-way ANOVA with Tukey’s post *hoc* test. *, *P* < 0.05; **, *P* < 0.01; ***, *P* < 0.001; ****, *P* < 0.0001; n.s., not significant.

### Polyfunctional T and B cells as well as memory responses in the lung and spleen stimulated by T4-CoV-Flu vaccines

In addition to humoral immunity, T-cell-mediated responses play a critical role in providing long-term protection against respiratory viruses^59,60^. To evaluate these responses elicited by the T4-CoV-Flu vaccines, we determined the phenotype and functionality of antigen-specific T cells in the spleen and lungs of i.n. immunized mice. Compared to T4 control group, intracellular cytokine staining revealed that T4-CoV-Flu (T4-M1-M2e-HA-S-ecto) i.n. vaccination led to a significant increase in the frequencies of S-ecto- and HA-stem-specific CD4^+^ and CD8^+^ T cells that produce IFNγ, IL-2, TNFα, and IL-17A in both spleen (Fig. 6, a to d) and lungs (Fig. 6, e to h). In contrast, the frequencies of antigen-specific T cells producing IL-4 and IL-5 did not significantly increase in all vaccinated groups except in mice immunized with T4-M1-M2e-HA-S that had higher splenic HA-specific CD4^+^ cells producing IL-4 and IL-5 (Fig. 6c). Most importantly, mice immunized with T4-CoV-Flu had overall higher cytokine positive T cells specifically for IL-2, TNFα and IL-17 in comparison with soluble trimer vaccinated mice (Fig. 6, a to h). These results indicated the T4-CoV-Flu-mediated induction of Th1 and cytotoxic T cell responses, which are desirable for COVID-19 vaccines, contributed to viral clearance and reduced disease severity^61,62^.

**Fig. 6:**
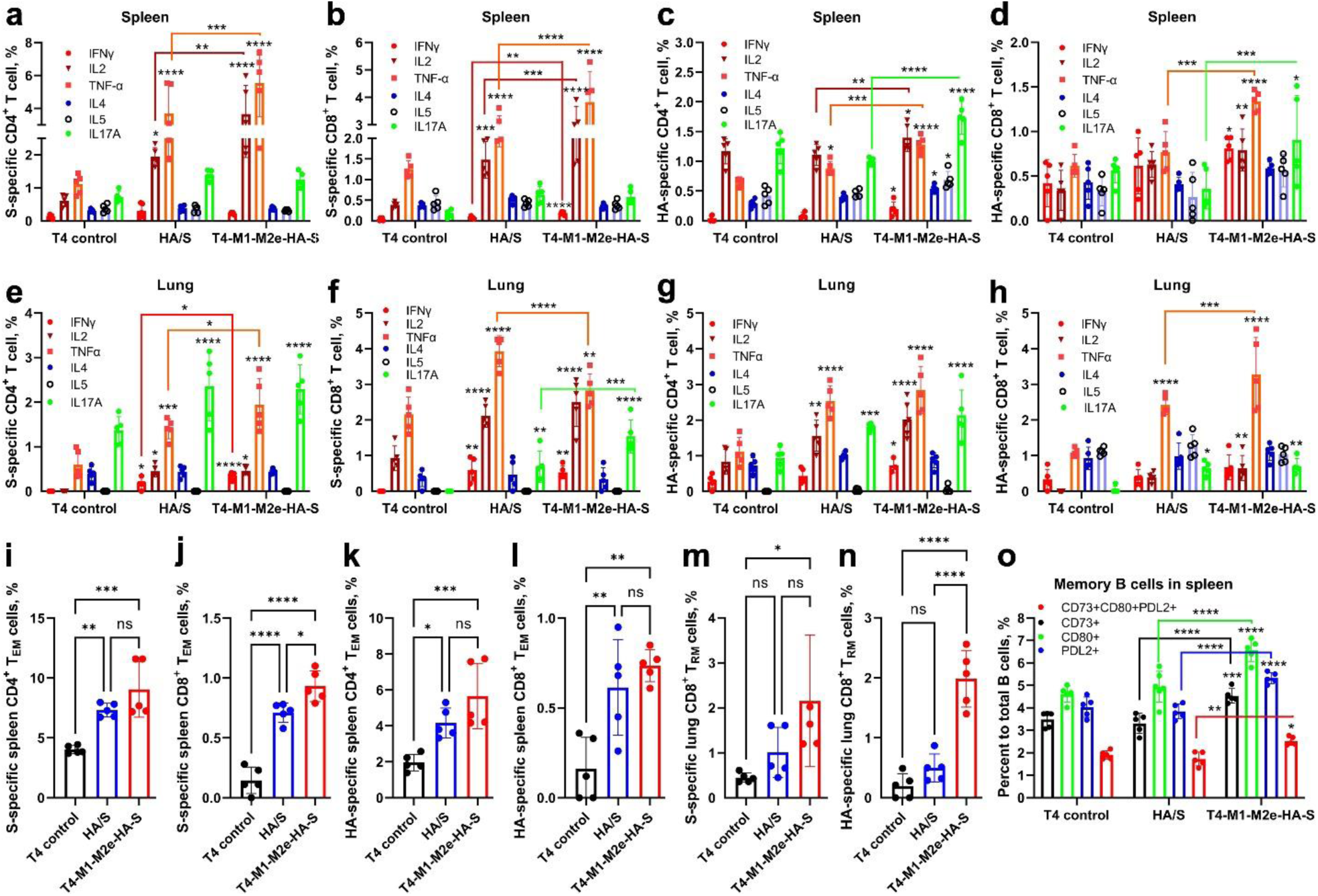
The T4-CoV-Flu intranasal vaccine elicits strong systemic and mucosal cellular immune responses. Mice immunized with T4 vector control, soluble HA-stem and S-ecto trimers (HA/S-trimer), or phage expressing HA-stem and S-ecto trimers (T4-M1-M2e-HA-S) were compared. Lungs and spleens were harvested after immunization and single-cell suspensions were prepared. Cells were stimulated with 100 µg/ml of either SARS-CoV-2 S-trimers or H1N1 HA-stem trimers for 16 hours, followed by an additional 4-hour incubation with Brefeldin A. Cells were then harvested and stained for various surface markers and cytokines. For T effector memory (T_EM_) cells: CD44^+^CD62L^-^CD127^-^. For lung CD8^+^ tissue-resident memory T cells (CD8^+^ T_RM_): CD8^+^CD4^-^CD44^+^CD69^+^CD103^+^CD49a^+^. For lung CD4^+^ T_RM_: CD4^+^CD8^-^CD44^+^CD69^+^CD11a^+^. For memory B cells, splenocytes were directly stained without stimulation for the phenotype: CD19^+^CD3^-^CD138^-^/CD38^+^GL7^-^/CD73^+^CD80^+^PDL2^+^. The percentages of these cells were plotted and compared among T4 control, soluble HA/S-trimer, and T4-M1-M2e-HA-S groups. **a** to **d** The percentages of S-ecto-specific CD4^+^ T cells (a) and CD8^+^ T cells (b), and HA-specific CD4^+^ T cells (c) and CD8^+^ T cells (d) secreting various cytokines (IFNγ, IL2, TNFα, IL4, IL5, and IL17A) in the splenocytes. **e** to **h** The percentages of S-ecto-specific CD4^+^ T cells (e) and CD8^+^ T cells (f), and HA-specific CD4^+^ T cells (g) and CD8^+^ T cells (h) secreting various cytokines (IFNγ, IL2, TNFα, IL4, IL5, and IL17A) in the lung. **i** to **l** The percentages of S-ecto-specific CD4^+^ T_EM_ cells (i) and CD8^+^ T_EM_ cells (j), and HA-specific CD4^+^ T_EM_ cells (k) and CD8^+^ T_EM_ cells (l) in the splenocytes. **m** and **n** The percentages of S-ecto-specific CD8^+^ T_RM_ (m) and HA-specific CD8^+^ T_RM_ (n) in the lung. **o** The percentages of memory B cells to total B cells in the splenocytes. Data represent means ± SD (n=5). Statistical comparisons among multiple groups were made using two-way ANOVA with Tukey’s post *hoc* test in (a to h) and (o). One-way ANOVA with Tukey’s multiple comparisons test was used to analyze statistical significance in (i to n). *, *P* < 0.05; **, *P* < 0.01; ***, *P* < 0.001; ****, *P* < 0.0001; ns, not significant.

Furthermore, T4-CoV-Flu vaccination led to a significant increase in the proportions of antigen-specific effector memory (T_EM_) CD8^+^ and CD4^+^ T cells in the spleen (Fig. 6, i to l), as well as tissue-resident memory (T_RM_) CD8^+^ T cells in the lungs (Fig. 6m, n). These memory T cell subsets are critical for providing rapid and effective recall responses upon re-exposure to the pathogen and are essential for maintaining long-term protective immunity^63^. However, S-ecto- and HA-stem-specific lung-resident CD4^+^ T_RM_ cells were not significantly increased in the T4-CoV-Flu vaccinated groups when compared to the T4 control (Supplementary Fig. 6a, b), suggesting that the T4-CoV-Flu vaccine may be more effective at inducing CD8^+^ T_RM_ cells than CD4^+^ T_RM_ cells in the lungs. The significant induction of lung-resident CD8^+^ T_RM_ cells by the T4-CoV-Flu vaccine is particularly promising, as these cells can provide immediate protection against respiratory viruses at the site of infection, even in the absence of circulating antibodies^64^.

The induction of memory B cells is crucial for maintaining long-term antibody production and providing rapid and enhanced antibody responses upon re-exposure to the pathogen^65^. The T4-CoV-Flu vaccine induced a significant increase in diverse memory B cells when compared to the soluble trimer vaccine (Fig. 6o; Supplementary Fig. 6c). Notably, the vaccine elicited diverse memory B cell populations, including CD73^+^, CD80^+^, PDL2^+^, and CD73^+^CD80^+^PDL2^+^ cells, all at higher levels than in control groups (Fig. 6o; Supplementary Fig. 6c). This combination of markers allows for the study of different functional aspects of memory B cells, such as immune regulation (CD73, PDL2) and activation (CD80). The diverse memory B cell response is consistent with reported studies showing that multi-antigen presentation on nanoparticles leads to greater B cell clonotype diversity and breadth of neutralization^20,21^. The enhanced diversity and quantity of memory B cells indicate the T4-CoV-Flu vaccine’s ability to elicit robust and potentially more broadly protective long-lasting adaptive immune responses.

Taken together, the above results demonstrated that i.n. vaccination with T4-CoV-Flu phages not only induced potent antibody responses, but also stimulated polyfunctional T cell responses in both systemic and mucosal compartments. Induction of antigen-specific CD4^+^ and CD8^+^ T cells producing Th1 and Th17 cytokines, diverse memory B cells, as well as generation of effector and lung tissue-resident memory CD8^+^ T cells, provided a compelling rationale for developing T4-CoV-Flu vaccine as a mucosal vaccine against multiple respiratory viruses.

### T4-CoV-Flu intranasal vaccine provided complete protection against lethal SARS-CoV-2 and influenza infections

To evaluate the protective efficacy of the T4-CoV-Flu vaccine, we challenged the immunized mice with lethal doses of either ancestral SARS-CoV-2 or influenza A/Puerto Rico/8/1934 (H1N1 PR8) virus and monitored their body weight and survival over time (Fig. 7a). In the SARS-CoV-2 challenge experiment, hACE2 mice vaccinated with T4 phage control rapidly lost weight and succumbed to the infection in 6 days. In contrast, mice immunized with T4-CoV (T4-NP-S) or T4-CoV-Flu (T4-M1-M2e-HA-S or T4-NP-HA-S) were completely protected against both weight loss and mortality (Fig. 7, b and c). Notably, the T4-CoV and T4-CoV-Flu vaccines provided superior protection compared to vaccination with soluble S-ecto/HA-stem trimers, which only conferred partial protection (60% survival rate) (Fig. 7, b and c), further highlighting the effectiveness of the T4-vectored mucosal delivery.

**Fig. 7:**
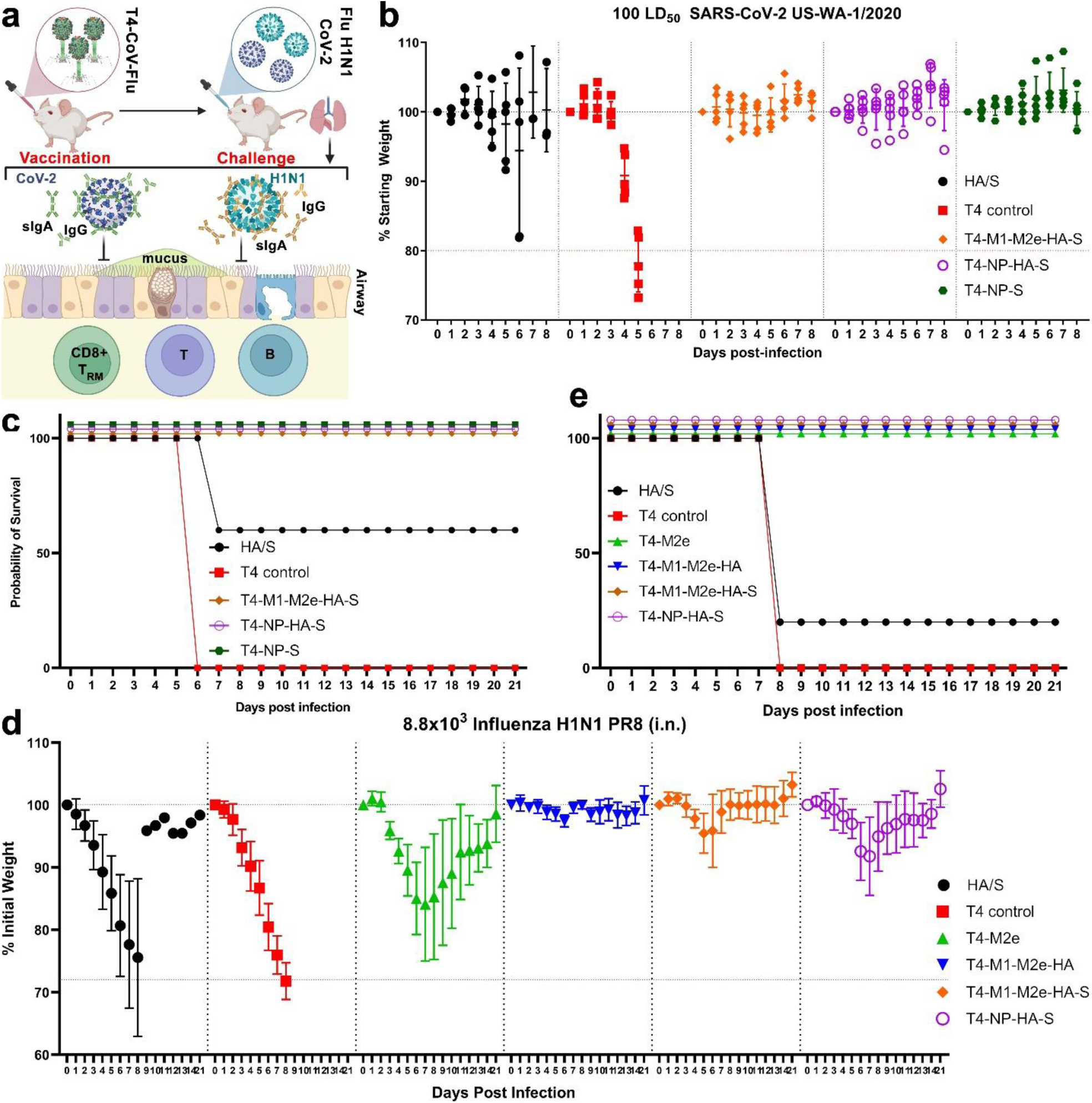
The T4-CoV-Flu intranasal vaccine conferred complete protection against high-dose SARS-CoV-2 and H1N1 challenges. **a** Schematic of T4-CoV-Flu intranasal vaccine protection against SARS-CoV-2 and influenza challenge through mouse airways. **b** and **c** Evaluation of the protective efficacy of T4-CoV-Flu vaccine candidates against SARS-CoV-2 in mice. (b) Percent starting body weight of immunized mice over days following intranasal challenge with 100 LD_50_ SARS-CoV-2 US-WA-1/2020. (c) Survival rate of mice following the SARS-CoV-2 challenge. **d** and **e** Assessment of the protective efficacy of T4-CoV-Flu vaccine candidates against influenza challenge. (d) Percent change in body weight of immunized mice over time after intranasal challenge with 8.8 x 10^3^ influenza H1N1 PR8. (e) Survival rate of mice following H1N1 PR8 challenge. The data are presented as means ± SD (n=5).

Similarly, in the influenza H1N1 challenge experiment, T4 control mice rapidly lost weight and succumbed to the infection in about 8 days. However, mice vaccinated with multivalent T4-Flu (T4-M1-M2e-HA) or T4-CoV-Flu (T4-M1-M2e-HA-S or T4-NP-HA-S) were completely protected against both weight loss and mortality. In contrast, soluble S-ecto/HA-stem trimers provided only low and partial protection, with a mere 20% survival rate (Fig. 7, d and e). Although the HA-specific serum antibody titers in the T4-M1-M2e-HA group were higher than the titers in the T4-M1-M2e-HA-S group (Fig. 4), there was no difference in body weight change between the two groups over 21 days following challenge (Supplementary Fig. 7a).

The co-display of HA-stem with other influenza antigens, such as M1 and M2e, in the T4-CoV-Flu vaccine (T4-M1-M2e-HA-S) contributed to improved protection, as evident by faster body weight recovery after challenge compared to the monovalent HA-stem antigen in the T4-NP-HA-S vaccine (Supplementary Fig. 7, b and c). Remarkably, T4 displaying M2e alone was able to confer complete protection against the H1N1 challenge, and incorporation of additional influenza antigens, M1 and HA stem, further significantly improved body weight recovery (Supplementary Fig. 7d). These findings supported the value of T4-arrayed multivalent display in enhancing the potency of vaccine-induced protection.

The above challenge studies demonstrated that nasal T4-CoV-Flu dual nanovaccines are effective in providing complete protection against lethal SARS-CoV-2 and influenza infections in mice. Thus, from the body of evidence described above, it can be concluded that the high-capacity antigen loading and plug-and-play nature of this platform, coupled with its ability to induce robust systemic and mucosal protective immunity, positions it as a promising needle-free vaccine delivery vehicle for combating respiratory pathogen pandemic and epidemic threats.

## Discussion

In this paper, we demonstrated a pinnacle VLP mucosal vaccine design by leveraging certain unique features of the noninfectious phage T4. The result is a multi-antigen, adjuvant-free, intranasal COVID-Flu dual vaccine that induces robust mucosal immunity and is highly efficacious against COVID-19 and Flu, two of the most widely circulating and deadly respiratory threats currently. Furthermore, our exhaustive studies address two of the most pressing concerns in vaccinology today, mucosal immunity and global vaccine engineering, for better pandemic preparedness against emerging bacterial and viral threats.

Our T4-CoV-Flu vaccine design exemplifies its remarkable adaptability for diverse antigen presentation, by incorporating five structurally and functionally distinct antigens from SARS-CoV-2 and influenza viruses in different compartments of a single nanoparticle. The final product carries arrays of approximately 56 copies of S-ecto trimers, 46 copies of HA-stem trimers, and 90 copies of M2e decorating the capsid as fusions of Soc and Hoc, while ∼40 copies of M1 or 100 copies of NP are encapsulated in the interior. Such a systematic and high-density epitope presentation and its ability to induce broad mucosal immunity, to our knowledge, has not been demonstrated in any other platform. Furthermore, the feasibility of rapidly and inexpensively producing phage up to 10^14^ -10^15^ particles per liter of *E. coli* highlights its potential for swift adaptation to evolving pandemic threats.

One of the most striking features of our vaccine is its ability to induce robust mucosal immunity, a critical factor in preventing respiratory virus transmission that is often lacking in injectable vaccines^11,18,66,67^. We observed elevated levels of sIgA in the BALF of vaccinated mice, far surpassing that induced by soluble trimers. This localized immune response, combined with strong systemic antibody production, provides a first line of defense as well as a dual layer of protection that could significantly reduce both infection and transmission rates.

The balanced Th1/Th2 immune response elicited by the T4-CoV-Flu vaccine is another key advantage over other vaccines. We observed high levels of IgG2a and IgG1 antibodies, with balanced ratios for both S-ecto and HA-specific responses. This balanced response is crucial for effective viral clearance and may reduce the risk of vaccine-enhanced respiratory diseases^54^. Furthermore, we demonstrated the induction of polyfunctional T cell responses, including a significant increase in the frequencies of antigen-specific CD4^+^ and CD8^+^ T cells producing IFNγ, IL-2, TNFα, and IL-17A in both spleen and lungs. Notably, we observed a marked increase in tissue-resident memory CD8^+^ T cells in the lungs of T4-CoV-Flu vaccinated mice suggesting potential for long-lasting local immunity^64^. Additionally, the vaccine induced diverse memory B cell populations with CD73, CD80, and PDL2 expression, suggesting enhanced immune regulation and activation potential that may contribute to broader, more durable protection against viral variants.

The complete protection observed in mice against lethal challenges of both SARS-CoV-2 and influenza viruses is a powerful demonstration of the vaccine’s efficacy. All mice vaccinated with T4-CoV or T4-CoV-Flu constructs survived the SARS-CoV-2 challenge without weight loss, compared to only partial survival in the soluble trimer group. Similarly, all vaccinated mice survived the influenza challenge versus a low survival rate in the soluble trimer group. This dual protection, achieved without the need for adjuvants, suggests that the intrinsic properties of the T4 nanoparticle may serve as a natural immune stimulant.

From a translational perspective, the T4-CoV-Flu vaccine offers several advantages that could significantly impact clinical practice and public health strategies. The intranasal administration route not only induces robust mucosal immunity but also offers the potential for needle-free vaccination, which could improve vaccine acceptance and simplify mass vaccination campaigns^68^. The stability of the T4 nanoparticle at room temperature^49^ could greatly facilitate vaccine distribution, particularly in resource-limited settings, by reducing cold chain requirements.

The modular nature of the T4 platform, allowing for the rapid incorporation of new antigens, positions this technology as a valuable tool for pandemic preparedness. We demonstrated that new recombinant phages could be generated in 2-3 weeks using our CRISPR-based engineering approach, which could dramatically reduce response times during future outbreaks. Moreover, the potential to combine antigens from multiple pathogens in a single vaccine could streamline immunization programs, improving coverage and reducing the logistical burden of multiple vaccinations.

However, several important questions and challenges remain to be addressed as we move towards clinical translation. While our results in mice are highly encouraging, studies in larger animals, particularly non-human primates, will be crucial to validate the vaccine’s efficacy in a model more closely resembling human physiology.

The mechanism by which the T4 nanoparticle enhances immune responses without traditional adjuvants warrants further investigation. The multivalent and symmetrized display of antigens on the T4 capsid may enhance B cell receptor cross-linking and recruitment of low-affinity B cells, leading to greater B cell clonotype diversity and breadth of neutralization^21–23^. Understanding these intrinsic immunostimulatory properties could provide valuable insights for the design of future mucosal nano-vaccine platforms.

Looking ahead, the evaluation of cross-protection against a broader range of SARS-CoV-2 variants and influenza strains will be critical to assess the breadth of protection. Studies on the duration of immunity, particularly the longevity of mucosal and tissue-resident memory responses, will inform vaccination strategies.

The global health implications of this technology are substantial. A single nasal vaccine protecting against both COVID-19 and influenza could significantly reduce the burden on healthcare systems, particularly during winter months when both viruses circulate. The potential for room-temperature stable, needle-free vaccines could revolutionize vaccination programs in low- and middle-income countries, contributing to greater global health equity.

In conclusion, our T4-CoV-Flu intranasal vaccine design represents a promising new approach in the fight against respiratory pathogens. Unlike many live or live-attenuated intranasal viral vaccine models currently under development, which can face challenges related to preexisting immunity and safety concerns, the noninfectious T4 platform provides a powerful alternative approach. By stockpiling a recombinant T4 phage scaffold containing *in vivo*-expressed and displayed antigens, any pandemic antigen(s) can be rapidly produced in bacterial, insect cell, or mammalian cell expression systems and loaded on the capsid by simple mixing of the two to create a variety of multivalent vaccines. Such protein-based T4 mucosal vaccines can be inexpensively and rapidly manufactured in response to emerging pandemic threats. Importantly, they can be mass-distributed without cold-chain requirements to resource-limited settings and to military personnel, even in remote areas of the world.

## Methods

### DNA, bacteria, bacteriophages, and viruses

The pET28b vector (Novagen, MA) was utilized to construct donor plasmids. Previously established methods^38,39,69^ were followed for constructing the LbCas12a and SpCas9 spacer plasmids. GeneArt (Thermo Fisher, MA) synthesized the donor DNA fragments that were pre-optimized with T4 codons for phage display. Plasmids containing the WT SARS-CoV-2 S-ecto-6P gene were generously provided by J. S. McLellan from the University of Texas, Austin. Plasmids with SpyCatcher/SpyTag sequences were obtained from Addgene (MA) (#133449).

For all cloning procedures, *E. coli* DH5α [hsdR17 (rK–mK^+^) sup2] (New England Biolabs, MA) was employed. The *E. coli* B40 (sup1) strain was used to propagate and produce wild-type and various recombinant phages.

The SARS-CoV-2 US-WA-1/2020 strain was obtained through CDC and is available at the World Reference Center for Emerging Viruses and Arboviruses, Galveston National Laboratory, UTMB. Influenza A virus (H1N1) strain A/PR/8/34 was propagated in specific pathogen-free chicken embryonated eggs and was purchased from the AVSBio (Norwich, CT). This strain was originally isolated in 1934 from a human patient in Puerto Rico.

### CRISPR engineering of T4 phage genome

The bacteriophage T4 genome was modified using a CRISPR-Cas-assisted editing approach as described previously with some modifications^38,55,69^. Briefly, guide RNAs for either the Cas9 nuclease from *Streptococcus pyogenes* or the Cas12a (Cpf1) nuclease from *Lachnospiraceae* bacterium were designed to target non-essential regions of the T4 genome selected for insertion of the SARS-CoV-2 and influenza viral genes. These regions included the Hoc, Soc, IPII, and IPIII loci (Fig. 1). The protospacer sequences were cloned into the respective CRISPR plasmids containing pCas9 or pCas12a nuclease gene, necessary trans-activating CRISPR RNA (tracrRNA), or direct repeat sequences. In parallel, donor plasmids were synthesized comprising the codon-optimized exogeneous genes flanked by ∼500 bp T4 homology arms matching the genomic insertion sites. The CRISPR and donor plasmids were co-transformed into chemically competent *E. coli* strains (DH5α or B40) and used to generate recombinant phages harboring the desired gene insertions following infection with wild-type T4 phage. Escaped phages that avoided cleavage by Cas9 or Cas12a due to successful incorporation of the donor sequence were enriched through a second round of infection in the CRISPR-expressing cells. Multiple iterative rounds of CRISPR-mediated editing and phage infection were performed to combine several modifications into a single phage genome, yielding the final multivalent T4-CoV, T4-Flu, or T4-CoV-Flu vaccine constructs. Genomic insertions were verified by PCR and Sanger sequencing. High-titer purified phage stocks were prepared by CsCl gradient ultracentrifugation.

### Phage production and purification

T4-CoV, T4-Flu, and T4-CoV-Flu phages were produced and purified as previously described^49,55^. Briefly, phages were propagated in *E. coli* strain B40 cultured in Moore’s medium. After infection and lysis, phages were harvested by centrifugation and purified using two rounds of CsCl gradient ultracentrifugation, followed by dialysis and 0.22-μm filtration. Phage concentration and antigen display were quantified by 4-20% SDS-PAGE.

### Expression and purification of prefusion-stabilized spike and HA stem trimers

Prefusion-stabilized ectodomain trimers of the SARS-CoV-2 spike protein (aa 1-1208 of the Wuhan-Hu-1 strain) with six proline substitutions was purified as described^49^. The influenza HA stem construct was designed based on the A/Puerto Rico/8/1934 (H1N1) virus. The HA stem sequence was genetically fused to a N-terminal 8× histidine tag and a C-terminal SpyTag003 peptide to enable covalent conjugation to SpyCatcher-displaying phages. The SpyTagged HA stem sequence was cloned into the pAAV vector and expressed by transient transfection into HEK293F cells using 293fectin reagent (Thermo Fisher). Trimeric HA-stem trimers were purified from the clarified culture supernatants by immobilized metal affinity chromatography (IMAC) using HisTrap FF columns (Cytiva, MA) followed by size exclusion chromatography (SEC) on a HiLoad 16/600 Superdex 200 (preparation grade) column (GE Healthcare, IL). The purified trimers were flash-frozen in liquid nitrogen and stored at -80°C.

### *In vitro* conjugation of spike and stem trimers to T4 phage scaffold

The display of Spytag-containing spike and HA stem trimers on the T4-SpyCatcher phage *in vitro* was assessed using a modified co-sedimentation technique^55,70^. In brief, phage particles were purified by two rounds of CsCl gradient centrifugation, filtered through a 0.22-μm filter, and sedimented at 34,000g for 30 minutes in Protein-LoBind Eppendorf tubes. The sedimented phages were then washed twice with sterile PBS buffer (pH 7.4) and resuspended in the same buffer. To remove any aggregates, S-ecto or HA-stem trimers were also sedimented at 34,000g for 15 minutes. Subsequently, T4-SpyCatcher phages were incubated with S-ecto and/or HA-stem trimers, either sequentially or simultaneously, at 4°C for 1-2 hours. The mixtures were centrifuged at 34,000g for 30 minutes to remove unbound proteins in the supernatants. The phage pellets were washed twice with excess PBS to eliminate any unbound protein and minor contaminants. The pellets containing the displayed proteins were then incubated at 4°C overnight and resuspended in PBS. The resuspended pellets were analyzed using a Novex 4-20% gradient SDS-PAGE mini gel (Thermo Fisher) to quantify the S-ecto or HA-stem trimer copies. After staining with Coomassie Blue R-250 (Bio-Rad, CA) and destaining, the protein bands on the SDS-PAGE gels were scanned and quantified using a ChemiDoc MP imaging system (Bio-Rad) and Image J software. The copy numbers of SpyCatcher and displayed spike or HA stem trimer molecules per capsid were calculated using gp18 (major tail sheath protein; 138 copies) or gp23 (major capsid protein; 930 copies) as internal controls, along with S-ecto or HA-stem trimer protein standards.

### Western Blot (WB) analysis

Following three freeze-thaw cycles and Benzonase treatment, purified phage particles were boiled in the SDS loading buffer for 10 minutes. They were then separated using 4-20% SDS-PAGE and transferred onto a polyvinylidene difluoride (PVDF) membrane (Bio-Rad). The membrane was processed for binding of NP- or Hoc-specific antibodies as described previously ^55^. The enhanced chemiluminescence (ECL; Bio-Rad) bands were visualized using the Bio-Rad Gel Doc XR+ System and analyzed using Image Lab software.

### Animal vaccination

Animal experiments adhered strictly to the guidelines set forth in the Guide for the Care and Use of Laboratory Animals by the National Institutes of Health. The protocols received approval from the Institutional Animal Care and Use Committee (IACUC) of both the Catholic University of America (Washington, DC; Office of Laboratory Animal Welfare assurance number A4431-01) and the University of Texas Medical Branch (UTMB, Galveston, TX; Office of Laboratory Animal Welfare assurance number A3314-01). SARS-CoV-2 virus infections were conducted within an animal biosafety level 3 (ABSL-3) containment facility while influenza virus infections were conducted within an animal biosafety level 2 (ABSL-2) containment facility, both at UTMB under approved protocols.

Female BALB/c mice (Jackson Laboratory, ME) or hACE2-transgenic AC70 mice (Taconic Biosciences, NY), aged 4-6 weeks, were randomly assigned to groups (5 animals per group) and allowed a 14-day acclimation period. For intranasal (i.n.) vaccinations, mice were anesthetized via inhalation of isoflurane. They were then administered with 2.5 x 10^11^ T4-CoV, T4-Flu, or T4-CoV-Flu phage particles in 50 µL of PBS (25 µL per nare). Two vaccine doses were given 3 weeks apart following a prime-boost regimen. Negative control groups received T4 backbone vector phage without antigens. A mixture of equal amounts of soluble HA-stem and S-ecto trimers served as the positive control. Blood samples were collected from each animal and the sera were stored at -80°C.

### SARS-CoV-2 and influenza H1N1 challenge

For animal challenge studies, mice were ear-tagged and weighed before the challenge to monitor individual weights throughout the study. Animals were anesthetized with isoflurane and then 50µL of viral challenge stock was administered (25 µL to each nare) followed by a 20µL PBS wash. For animals receiving SARS-CoV-2 US-WA-1/2020, the challenge dose was 100LD_50_ (3.3×10^2^ TCID_50_)^49,71^. For animals challenged with Influenza, the challenge dose was 8.8×10^3^ PFU (∼10LD_50_). Animals were weighed daily and scored for clinical signs of disease according to our approved IACUC protocols.

### Collection of Bronchoalveolar Lavage Fluid (BALF)

Around 21 days post influenza A PR8 strain challenge, survivals were euthanized, and BALF was collected using a method adapted from a prior study^72^. In brief, the tracheas were exposed by dissecting the salivary glands. A small cut was made on the trachea’s ventral side, and a blunt 26G needle, secured by tying the trachea around the catheter with floss, was inserted into the trachea. The lungs were then flushed with 600 μL of PBS using a 1-mL syringe, and the BALF samples were collected.

### Antibody ELISAs

To quantify SARS-CoV-2 spike, RBD, and NP-specific antibodies, as well as influenza HA and M2e-specific IgG, IgG1, IgG2a, and IgA antibodies in serum and BALF, we employed an enzyme-linked immunosorbent assay (ELISA). High-binding 96-well plates (Evergreen Scientific, CA) were coated overnight at 4°C with 100 µL of 1 µg/mL purified recombinant spike, RBD, NP, HA stem, or M2e peptide diluted in coating buffer (0.05 M sodium carbonate-sodium bicarbonate, pH 9.6). The plates were washed twice with PBS buffer, then blocked with 200 µL per well of PBS (pH 7.4) containing 5% BSA at 37°C for 2-3 hours. Serum and BALF samples were subjected to a 5-fold serial dilution, starting with an initial 100-fold dilution in PBS containing 1% BSA. Subsequently, 100 µL of the diluted samples were added to each well, and the plates were incubated at 37°C for 1 hour. The plates were then washed five times with PBST (PBS with 0.05% Tween 20). Following this, a secondary antibody, diluted 1:10,000 in PBS with 1% BSA (100 µL/well), was added. This included goat anti-mouse IgG-HRP, goat anti-mouse IgG1-HRP, goat anti-mouse IgG2a-HRP, or goat anti-mouse IgA-HRP (Thermo Fisher). After another hour of incubation at 37°C and five additional washes with PBST, the plates were developed using the TMB (3,3’,5,5’-tetramethylbenzidine) Microwell peroxidase substrate system (KPL, MA; 100 µL) for 5 to 10 minutes. The reaction was stopped by adding 100 µL of TMB BlueSTOP solution (KPL). The absorbance at 650 nm was read within 30 minutes using a VersaMax spectrophotometer. The endpoint titer was defined as the highest reciprocal dilution of serum or BALF that produced an absorbance greater than 2-fold the mean background of the assay.

### Foci Reduction Neutralization Test (FRNT)

*Cells and Viruses:* Vero E6 cells expressing human transmembrane serine protease 2 (TMPRSS2; JCRB1819) were obtained from the Japanese Collection of Research Bioresources Cell Bank (Ibaraki City, Osaka, Japan) and grown at 37°C with 5% CO_2_ in Dulbecco’s modified eagle medium (DMEM) (Gibco, NY) supplemented with 10% fetal bovine serum (FBS) (R&D Systems, GA) and 100 µg/ml Geneticin G418 (Sigma Aldrich, MO). The SARS-CoV-2 USA_WA1/2020 stock (lineage A) was passaged five times in Vero E6 cells and screened by next generation sequencing to confirm its identity as well as lack of tissue culture adaptations or mycoplasma contamination.

*Assay:* Antibody neutralization was assessed by FRNT essentially as previously described^73^. Mouse sera were heat-inactivated for 30 minutes at 56°C. Samples were serially 1:2 diluted in Dulbecco’s phosphate buffered saline (DPBS) (Corning, VA) over a range spanning 1:10 to 1:2,560. These dilutions were performed with independent triplicates. Control wells, which contained DPBS with no antibody, were included in triplicate on each plate. An equal volume of SARS-CoV-2, diluted in DPBS to a concentration of 5×10^3^ focus forming units (FFU/ml), was combined with the diluted sera such that the final antibody dilution factor range spanned 1:20 to 1:5,120. The virus and antibody were incubated for 1 hour at 37°C with 5% CO_2_ prior to 20 µL per well being transferred to 96-well plates with confluent monolayers of Vero E6 TMPRSS2 cells. The infection was allowed to proceed for 1 hour at 37°C prior to being overlaid with 85% minimum essential medium (Gibco) and 15% DMEM supplemented with 0.85% ^w^/_v_ methylcellulose (Sigma Aldrich), 2% FBS, 100µg/ml Geneticin G418, and 1% antibiotic-antimycotic (Sigma Aldrich).

The assays were incubated at 37°C with 5% CO_2_ for a further one day prior to fixation in 10% buffered formalin. Fixed monolayers were washed three times in DPBS before being incubated in permeabilization buffer (0.1% ^w^/v each of BSA [Sigma] and saponin [Sigma] in DPBS) for 30 minutes at room temperature. This buffer was replaced with permeabilization buffer containing a 1:1,000 dilution of polyclonal rabbit anti-SARS-CoV nucleocapsid antibody and allowed to incubate at 4°C overnight. Unbound primary antibody was removed by three washes in DPBS, and HRP-linked goat anti-rabbit IgG (Cell Signaling, MA) diluted 1:2,000 in permeabilization buffer was added and allowed to bind for 1 hour at room temperature. The unbound secondary antibody was removed by three washes in DPBS, and KPL TrueBlue peroxidase substrate (SeraCare, MA) was added. The reaction was allowed to proceed until foci were visualized, at which time the substrate was removed and the plates rinsed with water. Images of each well were captured using the Cytation7 Imaging Reader (BioTek, VT), and foci were counted manually.

The percent reduction in foci was calculated in comparison to the average of the three control wells within each plate. This data was fitted using the [Agonist] vs. response – Find EC anything function of Prism with the following constraints: the bottom limit was set to 0, the top limit was set to 100, and the F was set to either 50 or 80 to calculate the EC_50_ or EC_80_ values, respectively.

### Flow cytometry

Four weeks post-boost, both the immunized and the control hACE2-transgenic AC70 mice (n=5/group) were euthanized. Lungs from each mouse were infused with 10 ml PBS to remove residue blood. Spleen and lung tissues were then collected, and single-cell suspensions were made^74,75^. For B cell staining, about 2 million splenocytes from each spleen suspension were used. The cells were first blocked with anti-CD16/32 and then stained with Live/dead dye eFluor 780 and fluorescence-conjugated antibodies specific for memory B cells (Supplemental Table 1). For T cell staining, about 2 million splenocytes from both spleen and lung suspension were used. The cells were first seeded into a 96-well plate and stimulated with S-ecto or HA-stem trimers (60-100 µg/ml) for 16 hours at 37°C followed by incubation with Brefeldin A for an additional 4 hours. Cells were then harvested and stained with Live/dead dye eFlor780 and fluorescence-conjugated antibodies specific for various T cell surface markers (Supplemental Table 1). After washing with FACS buffer, cells were treated with eBioscience™ Foxp3 permeabilization buffer (Thermo Fisher) and stained for various cytokine markers (Supplemental Table 1). The differential cell populations were then acquired on flow cytometer (BD Symphony A5 SE, NJ) and analyzed with FlowJo software. The memory B cells and effector memory T cells from spleen tissue were identified as CD19^+^CD3^-^CD138^-^/CD38^+^GL7^-^/CD73^+^CD80^+^PDL2^+^ and CD3^+^/CD4^+^or CD8^+^/CD44^+^CD62L^-^ CD127^-^, respectively. While Lung CD4 and CD8 T_RM_ were gated as CD3^+^CD4^+^CD8^-^ CD44^+^CD69^+^CD11a+ or CD3^+^CD8^+^CD4^-^CD44^+^CD69^+^Cd103^+^CD49a^+^, respectively.

### Photo credit

Mouse and virus images (Figs. 4a and 7a) were created using BioRender (BioRender.com). T4 structure and schematics were created using ChimeraX. The figure data were organized using Photoshop CS6 (Adobe, CA).

### Statistical analysis

Statistical tests were performed using GraphPad Prism v9 software. For comparing multiple groups, one-way or two-way ANOVA with Tukey’s post-*hoc* tests for multiple comparisons were used. A nonparametric Student’s *t*-test was used to compare two groups. *P* values < 0.05 were considered significant. *, *P* < 0.05; **, *P* < 0.01; ***, *P* < 0.001; ****, *P* < 0.0001; ns, not significant. All data were presented as mean ± standard deviation (SD).

## Data availability

The authors declare that all data supporting the findings of this study are included in the manuscript and its Supplementary Information files.

## Acknowledgments

The authors thank Michell Brooke and Rachael Reyna for their help in performing FRNT. We thank Dr. Shinji Makino, UTMB, for providing polyclonal rabbit anti-SARS-CoV nucleocapsid antibodies for performing FRNT. This research was supported by NIAID/NIH supplement grant 3R01AI095366-07S1 (sub-award 1100992-100) and in part by NIAID/NIH grants AI081726 and AI175340, NIDA/NIH Avant Garde Award DP1DA060580, and National Science Foundation grant MCB-0923873 to V.B.R. AKC was funded through the Sealy-Institute for Human Infection and Immunity (IHII) grant, UTMB, to perform these studies.

## Author contributions

V.B.R., A.K.C., and J.Z. conceptualized and supervised the project. J.Z., J.S., H.B., S.J., X.W., E.K.H., P.B.K., and P.T. performed majority of experiments. K.S. isolated lung cells. K.S.P., J.A.P., and J.W. conducted FRNT assays. J.Z., V.B.R., J.S., and A.K.C analyzed and interpreted data and drafted the manuscript. All authors reviewed and edited the manuscript.

## Supplemental Figures and Tables

**Supplementary Fig. 1:**
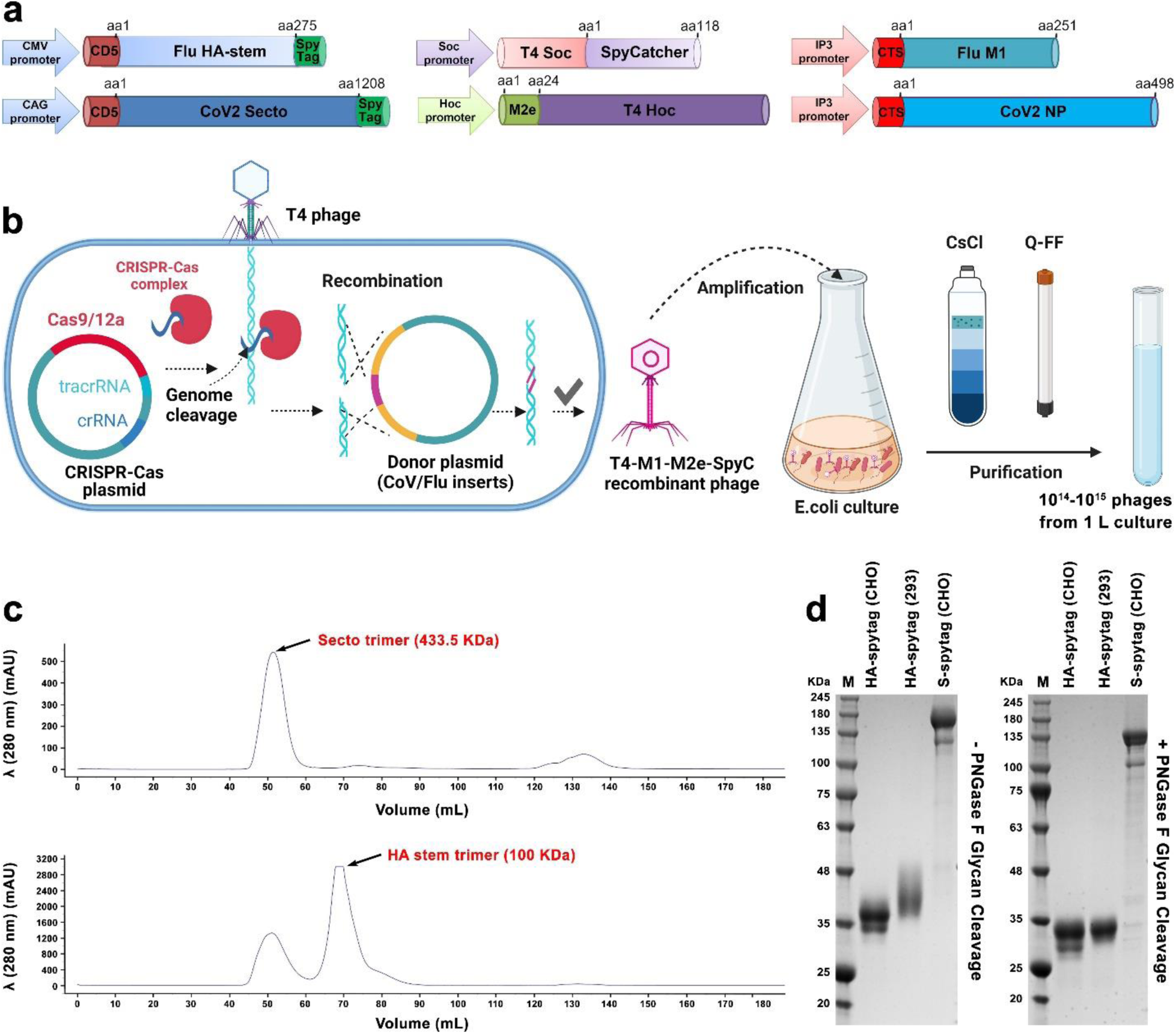
Components of the T4-CoV-Flu nanovaccine. **a** Schematic of various gene expression cassettes for T4 *in vitro* display, *in vivo* display, and encapsidation. **b** Schematic of CRISPR engineering scheme for the production of recombinant T4-M1-M2e-SpyCatcher phage. **c** and **d** Purification and characterization of S-ecto-Spytag trimers from ExpiCHO cells and HA stem-Spytag trimers from HEK293F cells. (c) Size-exclusion chromatography (SEC) elution profiles of S-ecto-Spytag trimers (top) and HA stem trimers (bottom). (d) SDS-PAGE patterns of SEC-purified HA stem-Spytag trimers (from ExpiCHO or HEK293F cells) and S-ecto-Spytag trimers (from ExpiCHO), both without (left) and with (right) the treatment of PNGase F glycan cleavage enzyme. Note the band shift of the envelope proteins following deglycosylation. The molecular weight standards (M) in KDa are shown on the left of each gel.

**Supplementary Fig. 2:**
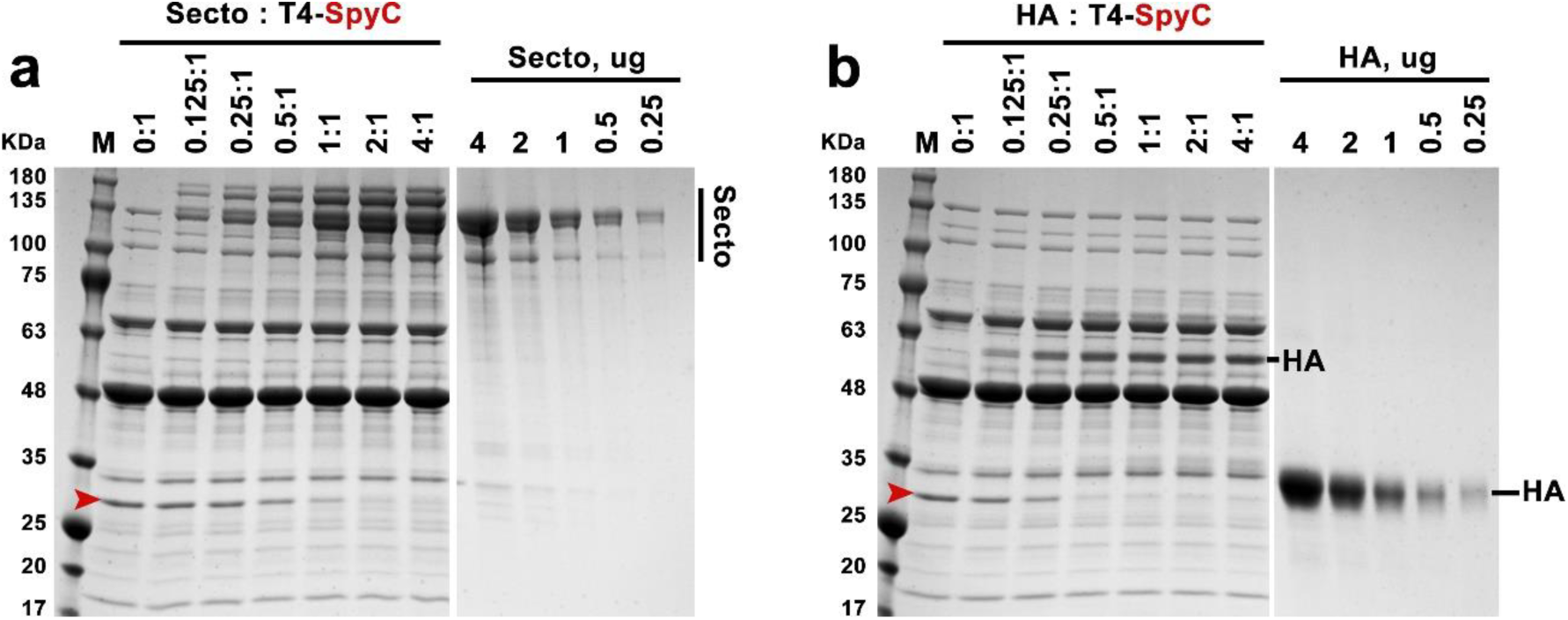
Optimization of S-ecto and HA-stem trimer display on T4 capsid. **a** *In vitro* display of S-ecto trimers on T4-SpyC phage at increasing ratios of S-ecto trimer molecules to Soc binding sites (0:1 to 4:1). S-ecto standard was used for quantification. **b** *In vitro* display of HA stem trimers on T4-SpyC phage at increasing ratios of HA stem trimer molecules to Soc binding sites (0:1 to 4:1). HA standard was used for quantification. Arrowheads depict the position of the Soc-SpyCatcher band which reduces in intensity as it gets conjugated with SpyTagged trimers with increasing SpyTagged trimers to Soc-SpyCatcher ratios.

**Supplementary Fig. 3:**
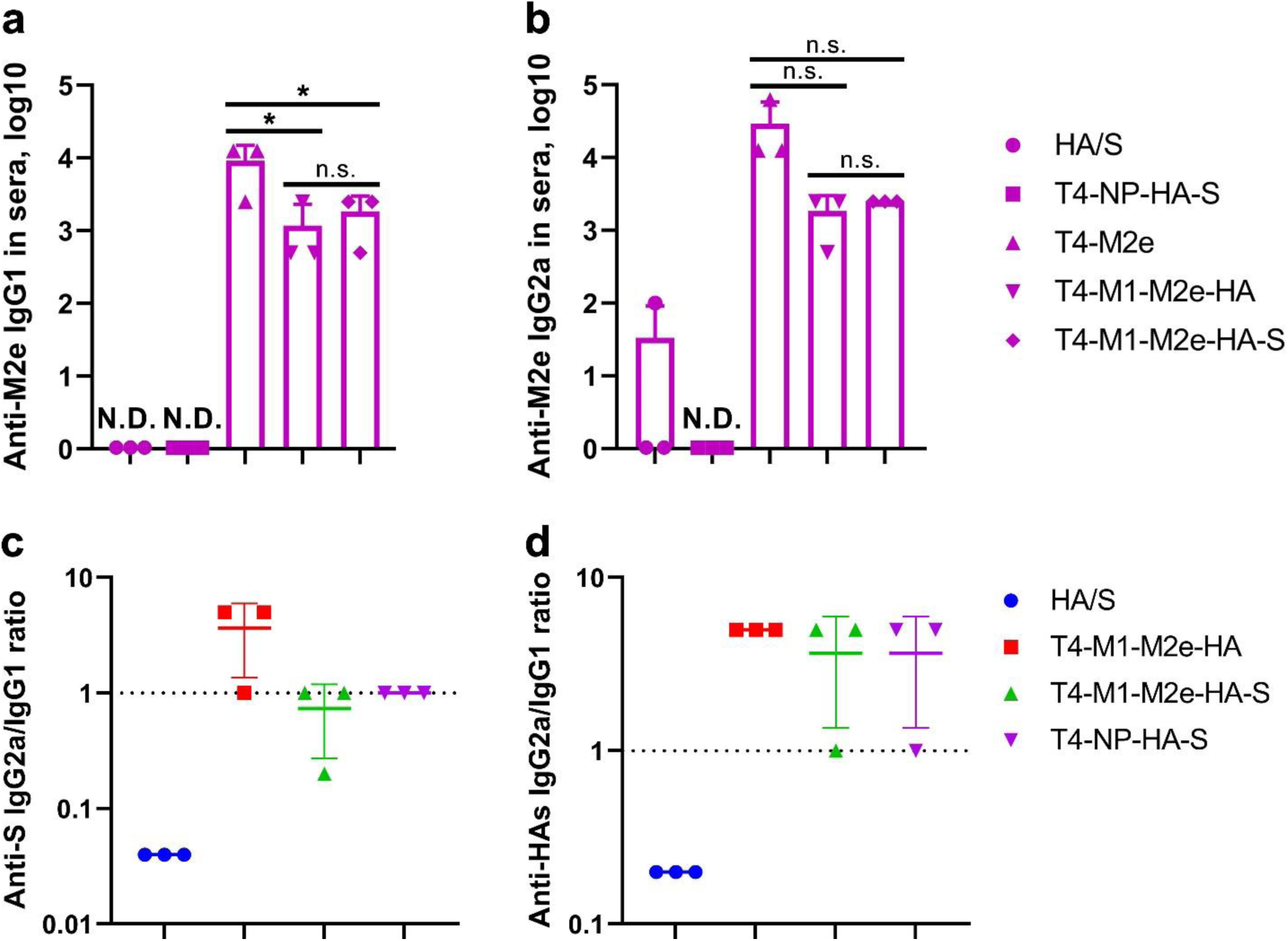
Intranasal immunization with T4-CoV-Flu vaccine elicited balanced systemic humoral immune responses. **a** and **b** Anti-M2e IgG1/IgG2a antibody responses in the sera of immunized BALB/c mice. **c** and **d** Comparison of IgG2a/IgG1 ratios for S-ecto-specific (c) and HA-stem-specific (d) antibodies in mouse sera across HA/S and various T4-CoV-Flu groups. The data are presented as means ± SD (n=3). Statistical comparisons among multiple groups were made using one-way analysis of variance (ANOVA) with Tukey’s post *hoc* test. *, *P* < 0.05; n.s., not significant.

**Supplementary Fig. 4:**
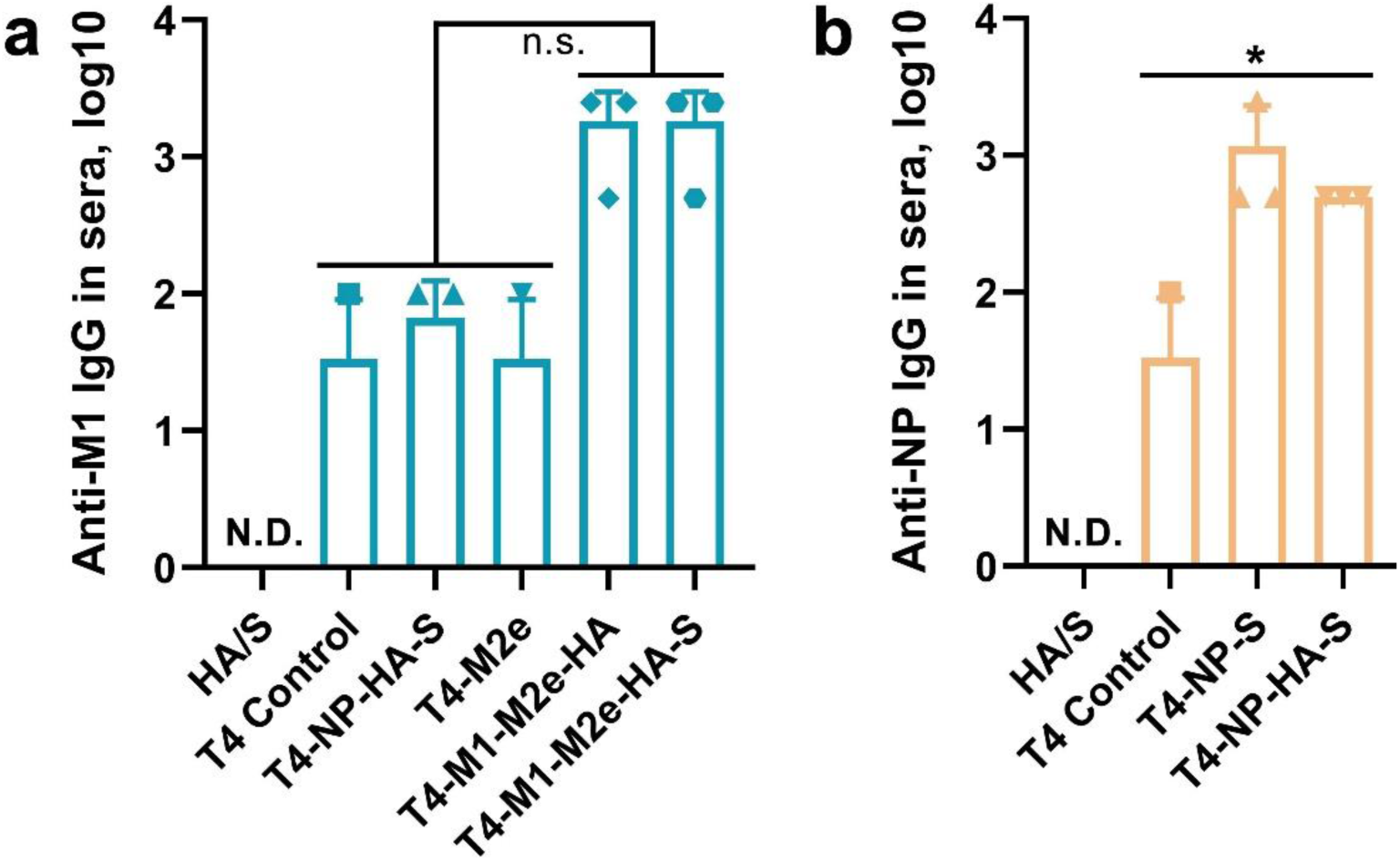
IgG antibody responses against M1 and NP in the sera of immunized mice. **a** Anti-Flu M1 IgG in BALB/c mice sera. **b** Anti-CoV NP IgG in ACE2 transgenic mice sera. ELISA was used to measure the reciprocal endpoint antibody titers. Data represent means ± SD. N.D. indicates not detected. The data are from three pooled independent experiments (n=5). Statistical comparisons among multiple groups were made using one-way ANOVA with Tukey’s post *hoc* test. *, *P* < 0.05; n.s., not significant.

**Supplementary Fig. 5:**
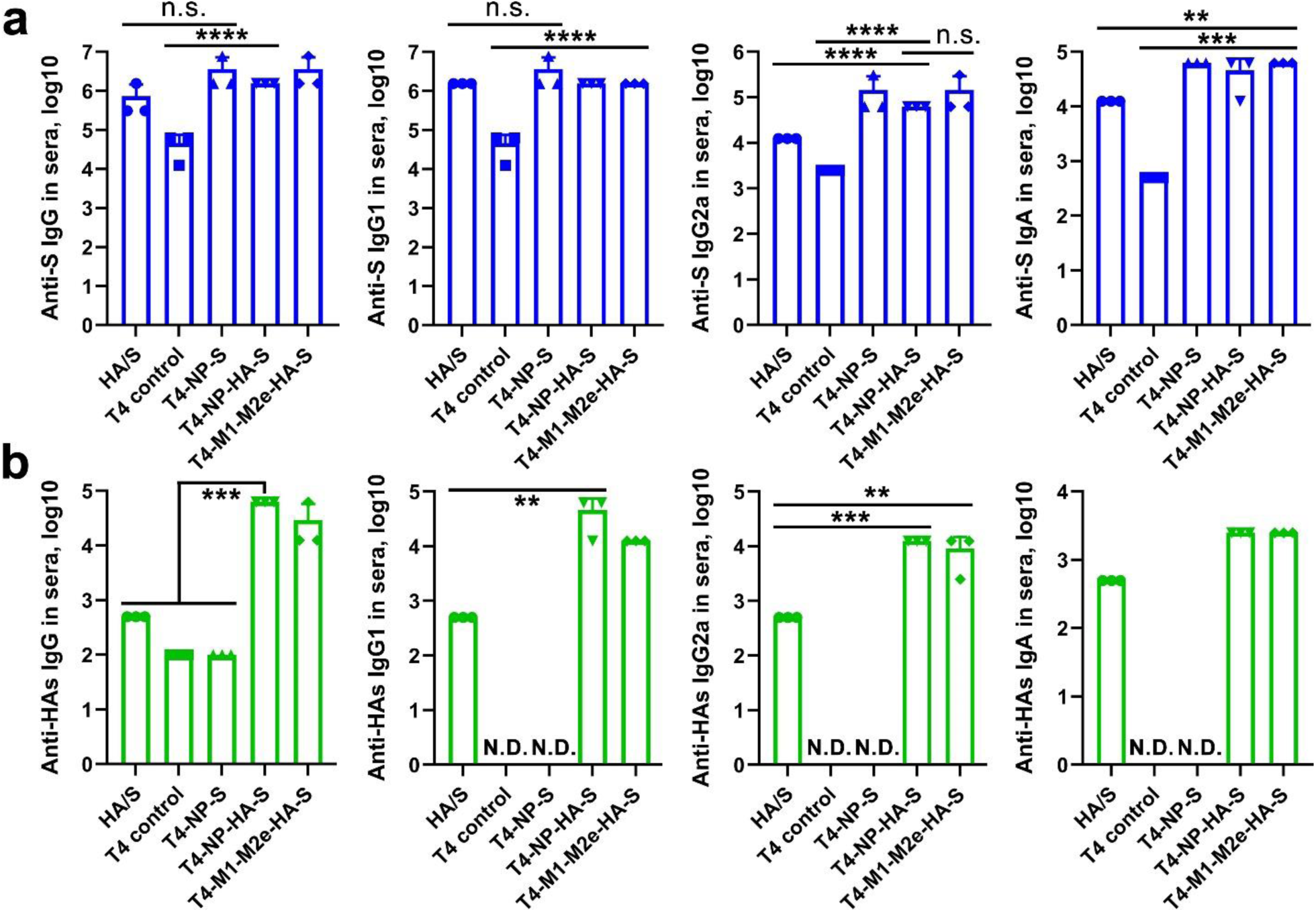
Antibody responses against CoV S-ecto and influenza HA-stem antigens in the sera of immunized hACE2 transgenic mice. ELISA was used to measure the reciprocal endpoint antibody titers for anti-S-ecto IgG/IgG1/IgG2a/IgA (**a**) and anti-HA-stem IgG/IgG1/IgG2a/IgA (**b**). Data represent means ± SD. N.D. indicates not detected. The data are from three pooled independent experiments (n=5). Statistical comparisons among multiple groups were made using one-way ANOVA with Tukey’s post *hoc* test. **, *P* < 0.01; ***, *P* < 0.001; ****, *P* < 0.0001; n.s., not significant.

**Supplementary Fig. 6:**
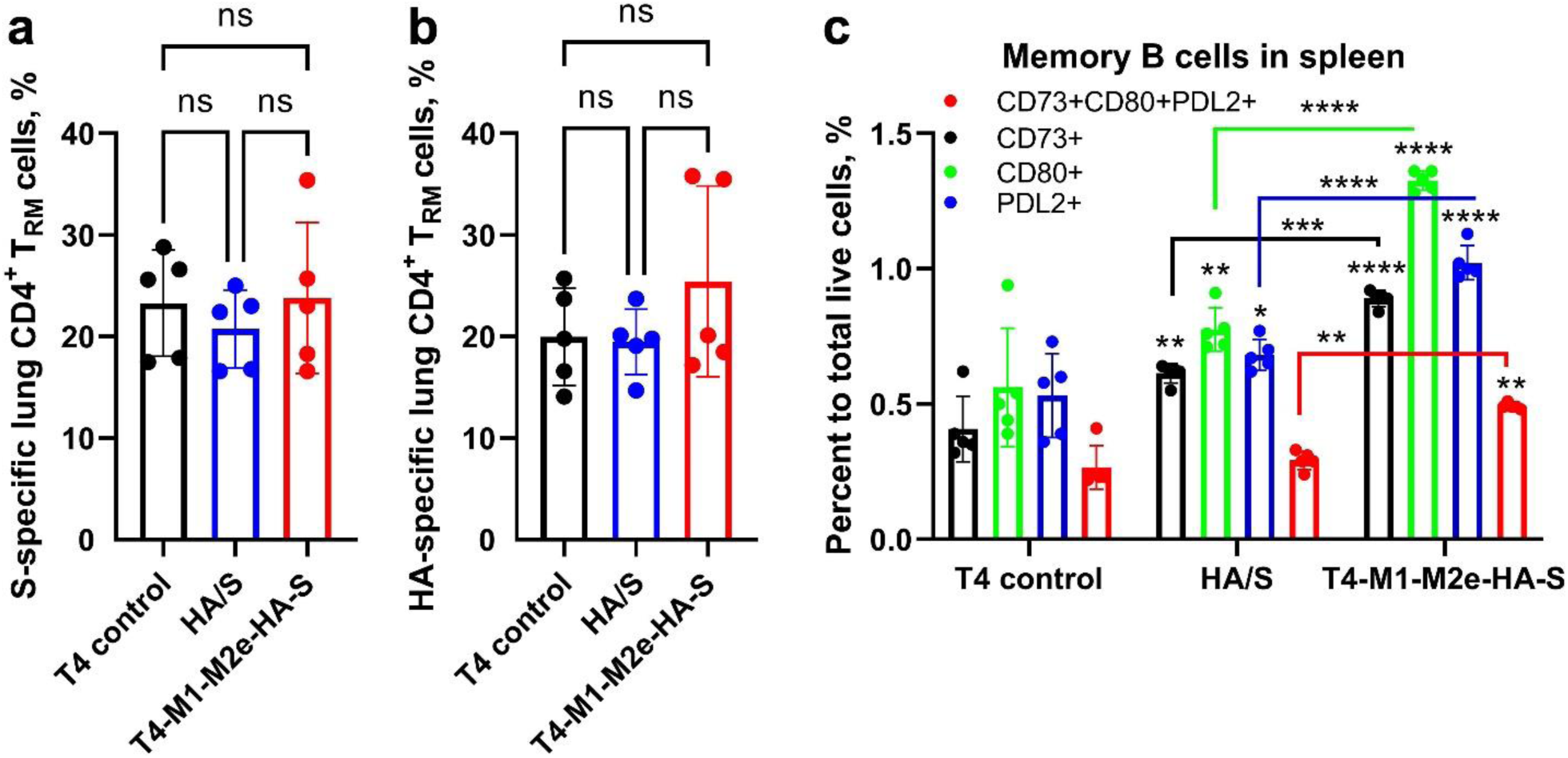
Comparison of memory B cells in spleen and CD4^+^ T_RM_ Cells in the lung among T4 control, soluble HA/S trimer, and T4-M1-M2e-HA-S groups. **a** and **b** The percentages of S-ecto-specific CD4^+^ T_RM_ (a) and HA-specific CD4^+^ T_RM_ (b) in the lung. **c** The percentages of memory B cells among total live cells in the splenocytes. Data represent means ± SD (n=5). One-way ANOVA with Tukey’s multiple comparisons test was used to analyze statistical significance in (a and b). Statistical comparisons among multiple groups were made using two-way ANOVA with Tukey’s post *hoc* test in (c). *, *P* < 0.05; **, *P* < 0.01; ***, *P* < 0.001; ****, *P* < 0.0001; ns, not significant.

**Supplementary S7:**
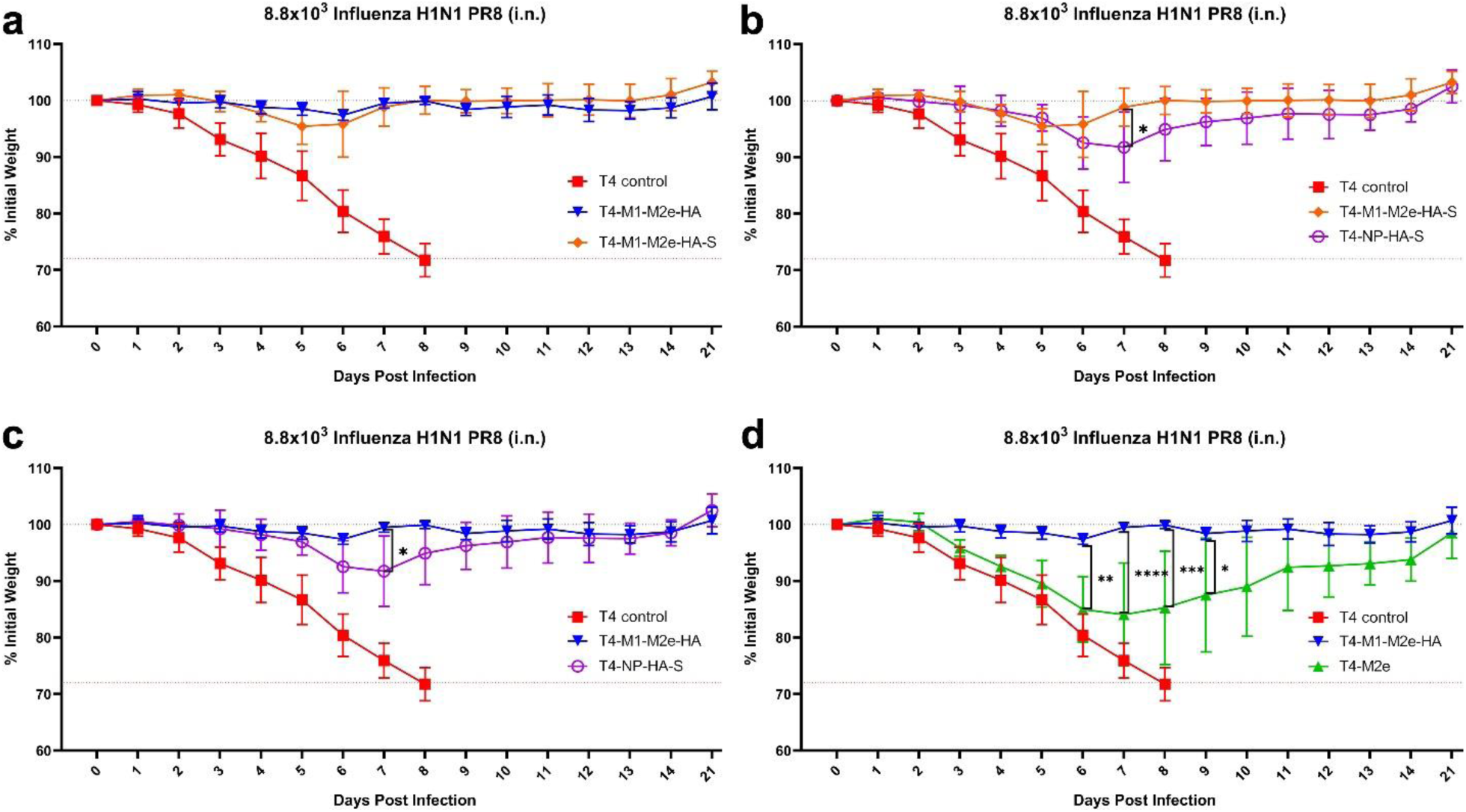
Body weight changes in T4-CoV-Flu intranasally-immunized mice following challenge with 8.8 x 10^3^ Influenza H1N1 PR8. Comparisons of body weights are shown: **a** Among T4 control, T4-MA-M2e-HA, and T4-M1-M2e-HA-S groups. **b** Among T4 control, T4-M1-M2e-HA-S, and T4-NP-HA-S groups. **c** Among T4 control, T4-M1-M2e-HA, and T4-NP-HA-S groups. **d** Among T4 control, T4-M1-M2e-HA, and T4-M2e groups. Data represent means ± SD (n=5). Statistical comparisons among multiple groups were made using two-way ANOVA with Tukey’s post *hoc* test. *, *P* < 0.05; **, *P* < 0.01; ***, *P* < 0.001; ****, *P* < 0.0001.

**Supplementary Table 1:** Antibodies used for Flowcytometry.

| Antibodies used for memory B cells |  |  |  |  |
| --- | --- | --- | --- | --- |
| Markers | Fluorophore | Clone | Vendor | Catalogue |
| CD19 | FITC | 6D5 | BioLegend | 115506 |
| CD3 | Alexa Fluor® 700 | 17A2 | BioLegend | 100216 |
| CD138 | BV650 | 281-2 | Beckton Dickinson | 564068 |
| CD38 | PE-Cy7 | 90 | BioLegend | 102718 |
| GL7 | Percp-cy5.5 | GL7 | BioLegend | 144610 |
| IgD | BV510 | 11-26c.a2 | BioLegend | 405723 |
| CD80 | BV421 | 16-10A1 | BioLegend | 104726 |
| CD73 | BV605 | TY/11.8 | BioLegend | 127215 |
| PDL2 | PE | TY25 | BioLegend | 107206 |
| Antibodies used for T cell surface staining |  |  |  |  |
| CD3 | Alexa Fluor® 700 | 17A2 | BioLegend | 100216 |
| CD4 | Brilliant Violet 785 | GK1.5 | BioLegend | 100453 |
| CD8 | FITC | 53-5.8 | BioLegend | 140403 |
| CD44 | BV510 | IM7 | BioLegend | 103044 |
| CD62L | BV711 | MEL-14 | BioLegend | 104445 |
| CD127 | PE | S18006K | BioLegend | 158204 |
| CD69 | BUV737 | H1.2F3 | BD Biosciences | 612793 |
| CD103 | BUV395 | M290 | BD Biosciences | 740238 |
| CD49a | BV750 | Hα31/8 | BD Biosciences | 746854 |
| CD11a | BUV805 | 2D7 | BD Biosciences | 741919 |
| Antibodies used for T cell intracellular staining |  |  |  |  |
| IFN-γ | PerCP/Cyanine5.5 | XMG1.2 | Biolegend | 505822 |
| IL-2 | PE-CF594 | JES6-5H4 | BD Biosciences | 562483 |
| TNF-α | eFluor 450 | MP6-XT22 | Thermo Fisher | 48-7321-82 |
| IL-4 | BV650 | 11B11 | BD Biosciences | 564004 |
| IL-5 | APC | TRFK5 | BD Biosciences | 554396 |
| IL-17A | PE/Cyanine7 | TC11-18H10.1 | Biolegend | 506922 |

